# Characterization of gut microbiomes in rural Honduras reveals novel species and associations with human genetic variation

**DOI:** 10.1101/2025.03.25.645309

**Authors:** Francesco Beghini, Ilana L. Brito, Mark Gerstein, Nicholas A. Christakis

**Affiliations:** Human Nature Lab, Yale University, New Haven, CT, USA; Meinig School of Biomedical Engineering, Cornell University, Ithaca, NY, USA; Program in Computational Biology and Biomedical Informatics, Yale University, New Haven, CT, USA; Department of Molecular Biophysics and Biochemistry, Yale University, New Haven, CT, USA; Department of Computer Science, Yale University, New Haven, CT, USA; Department of Biomedical Informatics and Data Science, Yale University, New Haven, CT, USA; Department of Statistics and Data Science, Yale University, New Haven, CT, USA

## Abstract

The gut microbiome is integral to human health, yet research data to date has emphasized industrialized populations. Here, we performed large-scale shotgun metagenomic sequencing on 1,889 individuals from rural Honduras, providing the most comprehensive microbiome dataset from Central America. We identify a distinct microbial composition enriched in *Prevotella* species, with 861 previously unreported bacterial species. Functional profiling reveals unique carbohydrate metabolism adaptations consistent with high-fiber diets. Longitudinal analysis over two years reveals microbiome instability, with shifts in taxonomic diversity and metabolic potential, including changes associated with SARS-CoV-2 infection. Additionally, we characterize the gut virome and eukaryotic microbiome, identifying novel viral taxa, including *Crassvirales* phages, and a high prevalence of *Blastocystis* species in individuals with greater microbial diversity. Finally, by integrating host genomic data obtained from low-pass saliva whole-genome sequencing, we uncover significant host-microbiome associations, highlighting the influence of human genetic variation on microbial composition. People who are more genetically similar also have more similar gut microbiomes. These findings expand our understanding of microbiome diversity in non-industrialized populations, highlighting the uniqueness of those microbiomes and underscoring the need for global microbiome research.

## Introduction

Extensive work has investigated the role that the human microbiome plays in health and illness, and how variations in diet, lifestyle, social interactions, and environment are relevant to its composition ^1–3^. At present, existing sampled microbiomes are heavily skewed towards western and industrialized countries in which individuals have generous access to antibiotics and other medications and a diet rich in ultra-processed food and low in fiber ^4,5^. Yet, more than 80% of the world population lives outside Europe and Northern America. An exploration of people’s microbiome composition from geographically diverse areas is necessary for a more comprehensive understanding of variation in microbiome composition, as well as its association with human health and even with host genetic background. Previous studies have documented that non-western populations harbor a more diverse microbiome, with several taxa that do not occur in other populations ^6,7^; in fact, some microbes appear to even have become extinct in western populations ^8^.

Modern tools have simplified the process of exploring this variation. Advances in sequencing and computational methods have enabled the reconstruction of more refined metagenomic assembled genomes (MAG), and the consequent expansion of the set of observed species in the human gut. MAGs reconstructed from under-represented regions can provide novel ecological and functional insights into the human diversity of gut microbiome.

Here, we perform shotgun metagenomic sequencing on 1,889 individuals living in the isolated western highlands of Honduras, which is the largest shotgun metagenomic data collected in the Central America region and from a single non-western population to date. In addition, using saliva, we performed low-pass sequencing of the human genome in order to conduct microbiome-genome association analyses. By putting the Honduras microbiome in a more global context (using previously collected data obtained by other investigators), we describe the extent of microbial diversity observed in Honduras and report several uncharacterized species obtained from metagenomic assembly and binning. We also explore associations between host human genotypes and microbiome occurrence as assessed by the microbiome genotypes.

## Results

### Gut microbiome genome catalog

We performed shotgun metagenomics on a total of 2,194 samples collected from 1,889 individuals living in 18 isolated villages in Honduras, which generated a total of 27.5 Tbp of data (mean 12.53 Gbp per person, stdev 4.4 Gbp). Of these, 301 individuals were sampled twice at a two-year interval, first in 2020 and then again in 2022.

We then proceeded to include our cohort in a larger context by integrating our analyzed sample set with human gut metagenomes from 29 different datasets spanning different lifestyles and geographical locations, resulting in a total of 10,173 metagenomes ^9–37^ (Figure S1, Table S1).

SGB-level (species-level genome bins)^21^ composition was determined with MetaPhlAn 4 ^38^. Similar to what has been previously observed in other non-western populations^8,30,39^, the overall species-level composition in the Honduras dataset is dominated by *Prevotella copri* clades, now reclassified as the *Segatella* genus; in particular, 52% of the samples have *Prevotella copri* clade A as the most abundant species, with a decreasing fraction of samples showing *Prevotella hominis* and SGB 3548 (*Spirochaetia* bacterium) as the most abundant species. We also observed species like SGB5809 (Veillonellaceae GGB4266), SGB1657 (Prevotellaceae GGB1239), and *Methanobrevibacter smithii* (SGB group 714) (Table S2) that were found to be more abundant, on average, in our cohort than other non-western cohorts. *Lacrimispora amygdalyna* (SGB 4716) was identified as the most prevalent species among our Honduras samples, present in virtually all the samples (99.5%).

### Comparison to other cohorts

While we did observe some overlap in composition between individuals in Honduras and African and Middle Eastern cohorts, the Honduras abundance profiles are nevertheless distinguishable (PERMANOVA *P*=0.001) (Figure 1A).

**Figure 1:**
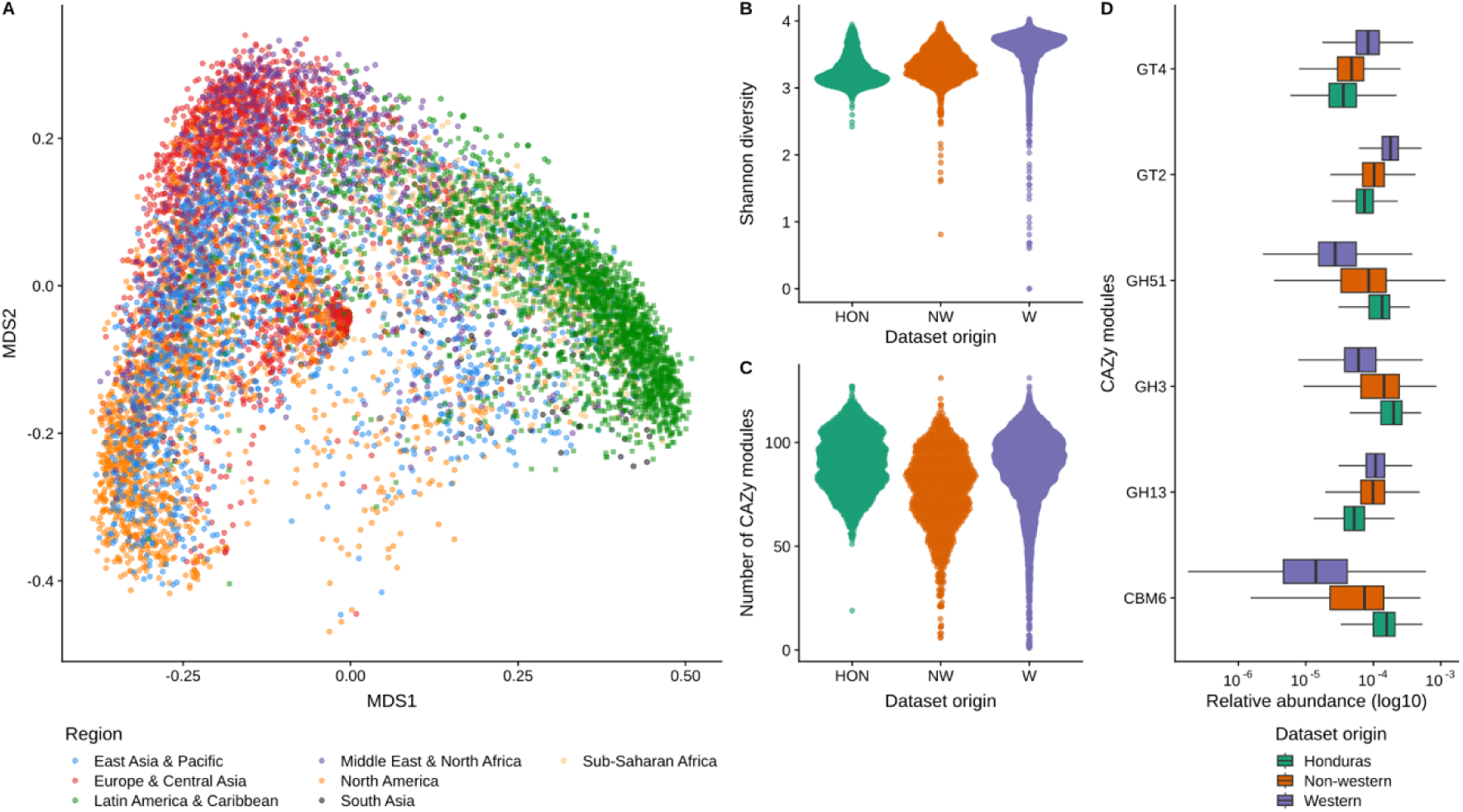
Principal Coordinate Analysis (A) of the Bray-Curtis dissimilarity calculated on the SGB’s relative abundance of 10,173 samples from curatedMetagenomicData including the 1,889 individuals in this study. Samples from Honduras (annotated with a square shape) overlap with other non-Western cohorts. Functional annotation with CAZy modules showed that Honduras samples have less diversity (B) but more richness (C), as well as having certain modules present in higher abundance compared to other datasets (D).

The same distinction can be observed as well in the functional composition, although this is not as clear as the taxonomic one (Figure S2). This is apparent when looking at the distribution of carbohydrate-active enzymes (CAZy), which digest carbohydrates; Principal Coordinate Analysis showed a clear separation between our cohort and other western and non-western cohorts. Differential abundances analysis showed that three CAZy classes – GH (Glycoside Hydrolase), GT (Glycosyl Transferases), and CBM (Carbohydrate Binding Module) – were present at different levels in our cohort. GH modules are enzymes that are able to break down carbohydrates into simpler sugars, and their presence is associated with diets high in fiber ^40^; CBMs are non-catalytic proteins able to enhance the catalytic activity of the enzymatic modules; and GT modules are glycosyltransferases. Higher median abundance of CBM6, GH3, GH51 modules and a lower median abundance of GH13, GT2, GT4 modules were observed in Honduras compared to other cohorts (Figure 1D).

Analysis of functional annotations of the reconstructed MAGs showed that those modules were present in higher prevalence in genera like *Prevotella*, *Bacteroides*, *Phocaeicola*, and *Butyribacter*. In contrast to previous reports ^8^, we observed that non-western cohorts exhibited lower richness of CAZy modules compared to western cohorts, while Honduras showed higher richness than both western and non-western cohorts (Figure 1B-C).

We compared the overall distribution of specific bacterial families to assess differences in composition between our cohort and other western and non-western cohorts. Families *Spirochaetaceae*, *Prevotellaceae*, and *Succinivibrionaceae* are commonly found to be increased in populations with traditional lifestyles (VANISH taxa), while industrialized population have higher abundances of *Verrucomicrobiaceae, Akkermansiaceae,* and *Bacteroidaceae* (BloSSUM taxa) ^7,8^ (Figure S3).

We identified a total of 170 species from the VANISH families and 185 species from the BloSSUM species. The overall distribution of BloSSUM taxa in the Honduras population resembles the one observed in other non-western cohorts, with slighter higher abundances of *Bacteroidaceae* taxa (Wilcoxon Test *P*=8.09 ×10^−22^), and lower abundances of *Verrucomicrobiaceae* than other non-western cohorts (Wilcoxon Test *P*=3.75×10^−14^). All three VANISH families are present in higher abundances, especially *Prevotellaceae* (Wilcoxon Test *P*=8.17×10^−18^). *Spirochaetaceae* and *Succinivibrionaceae* are found in lower abundances, but higher than other cohorts (*Spirochaetaceae* Wilcoxon Test *P*=4.73×10^−30^; *Succinivibrionaceae* Wilcoxon Test *P*=3.1×10^−2^)

In addition, our cohort had richer and more diverse samples (as measured by both the Shannon diversity and the number of observed species) than other analyzed countries (P<0.05) except for datasets from Ethiopia, Indonesia, Cameroon, Fiji, Indonesia, and Liberia (Figure S4).

We did not observe any differences in alpha diversity between people living in different villages in Honduras, but some villages could be distinguished according to differences in overall composition (PERMANOVA *P* = 0.001). Although most people self-reported their ethnicity as Maya Ch’orti’, we did not observe any difference in microbiome composition based on self-reported ethnicity.

### Strain-Level Analysis

To expand the understanding of intra-species diversity in our cohort, we investigated the strain-level diversity distribution by performing an analysis with StrainPhlAn. When considering the species most prevalent in our dataset, we found a wide range of strain diversity compared to the same species present in other datasets (Figure 2A, Table S3). *Haemophilus parainfluenzae* and *Enterobacter hormaechei* were the species with higher variation, with more than 3% of polymorphic sites, and hence a median number of polymorphic rates from Honduras strains higher then strains from other cohorts (Supplementary Table 3). This indicates higher intra-population strain diversity within Honduras was comparable to other sequenced cohorts.

**Figure 2:**
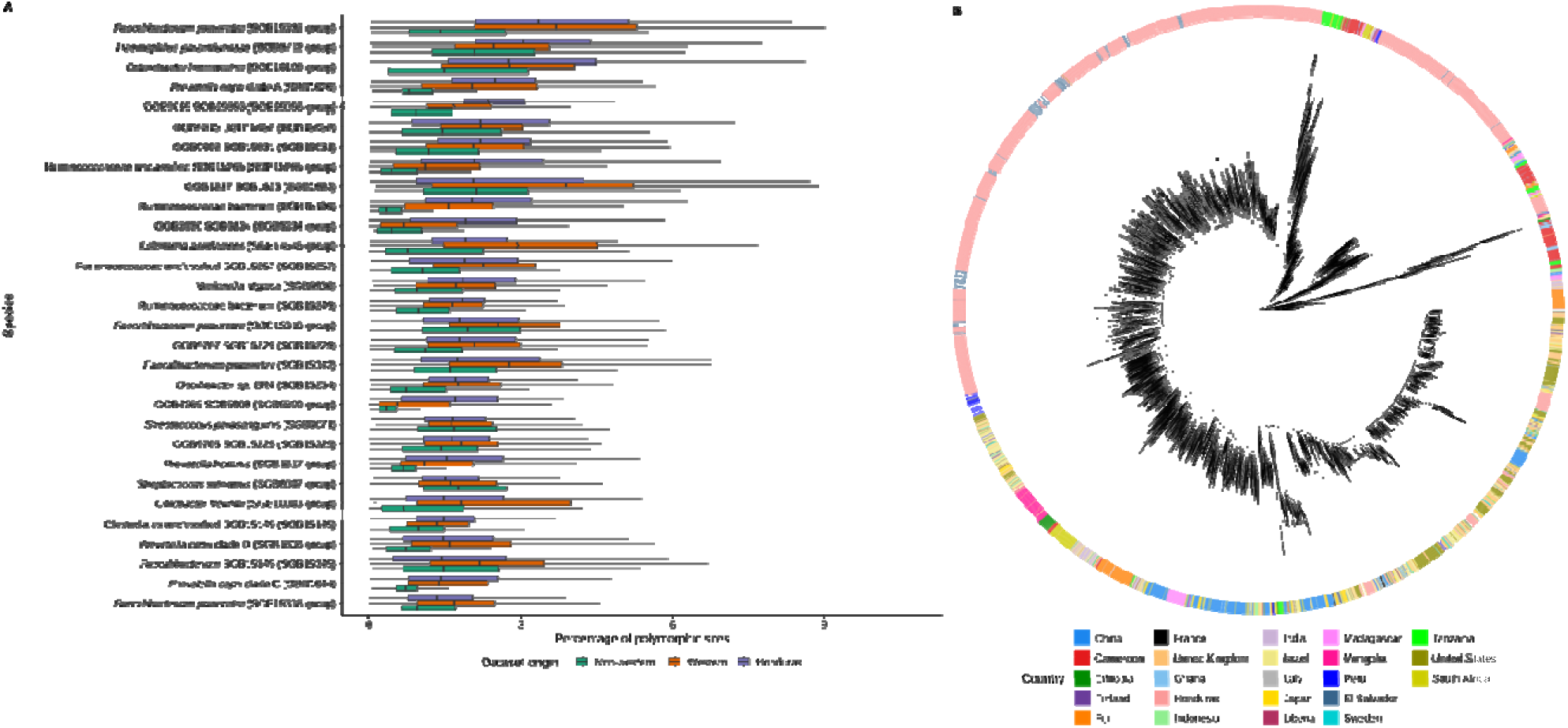
(A) Distribution of the percentage of polymorphic sites for 30 species identified by StrainPhlAn 4 (B) Phylogenetic tree of 5,001 *Prevotella copri* clade A strains reconstructed by StrainPhlAn 4. The sample country of origin is reported as a color annotation in the outer ring, strains reconstructed from Honduras samples (in pink) are segregated in branches significant associated with Honduras origin (D statistic: −0.40)

One species in particular, the butyrate-producing *Butyrivibrio crossotus,* presented high genetic (strain-level) diversity in our Honduras sample. *B. crossotus* is a component of the healthy gut microbiome; its prevalence is higher in samples from non-industrialized settings, and the species encodes several carbohydrate-active enzymes able to digest fiber^41,42^. We identified *B. crossotus* in nearly 80% of Honduras samples (n=1,729), with relative abundances as high as 15% in some samples. In other analyzed datasets, *B. crossotus* was found present in higher prevalence in non-industrialized cohorts and in lower prevalence in industrialized settings ^42^. *Prevotella copri* clade A, the most abundant species in our dataset, showed high intra-species diversity as well, with samples split into two primary separate and diverging clades, suggesting high geographical adaptation or the presence of another *P. copri* clade ^39^ not detected by the markers present in the reference database (Figure 2B).

Phylogenetic analysis of the *P. copri* strains retrieved from the Honduras samples revealed the presence of clades enriched in samples from a common origin, also previously reported by other studies ^43,44^. Honduras strains are segregated in branches that rarely overlap with samples with other geographical origins, with the exception of very few strains retrieved from El Salvador, a neighboring country; that is, we observed a significant association with Honduras origin (D statistic: −0.40, Predictive performance score (ELPD) −6149.7 ± 122.3). The same close association with geography was observed for SGB 1614, now identified as *Prevotella copri* clade H, in which multiple monophyletic groups are composed by strains from identical geographic locations, mostly from non-westernized populations. In addition, we observed the presence of a sub-species branch that is particularly divergent, with all strains retrieved from Honduras (Figure S5). This same pattern of phylogenetic clades enriched in samples retrieved from Honduras was observed in multiple species (not just *P. copri*), revealing strong geographic association of clades (n=1,545 species with ELPD < −100).

**Figure 3:**
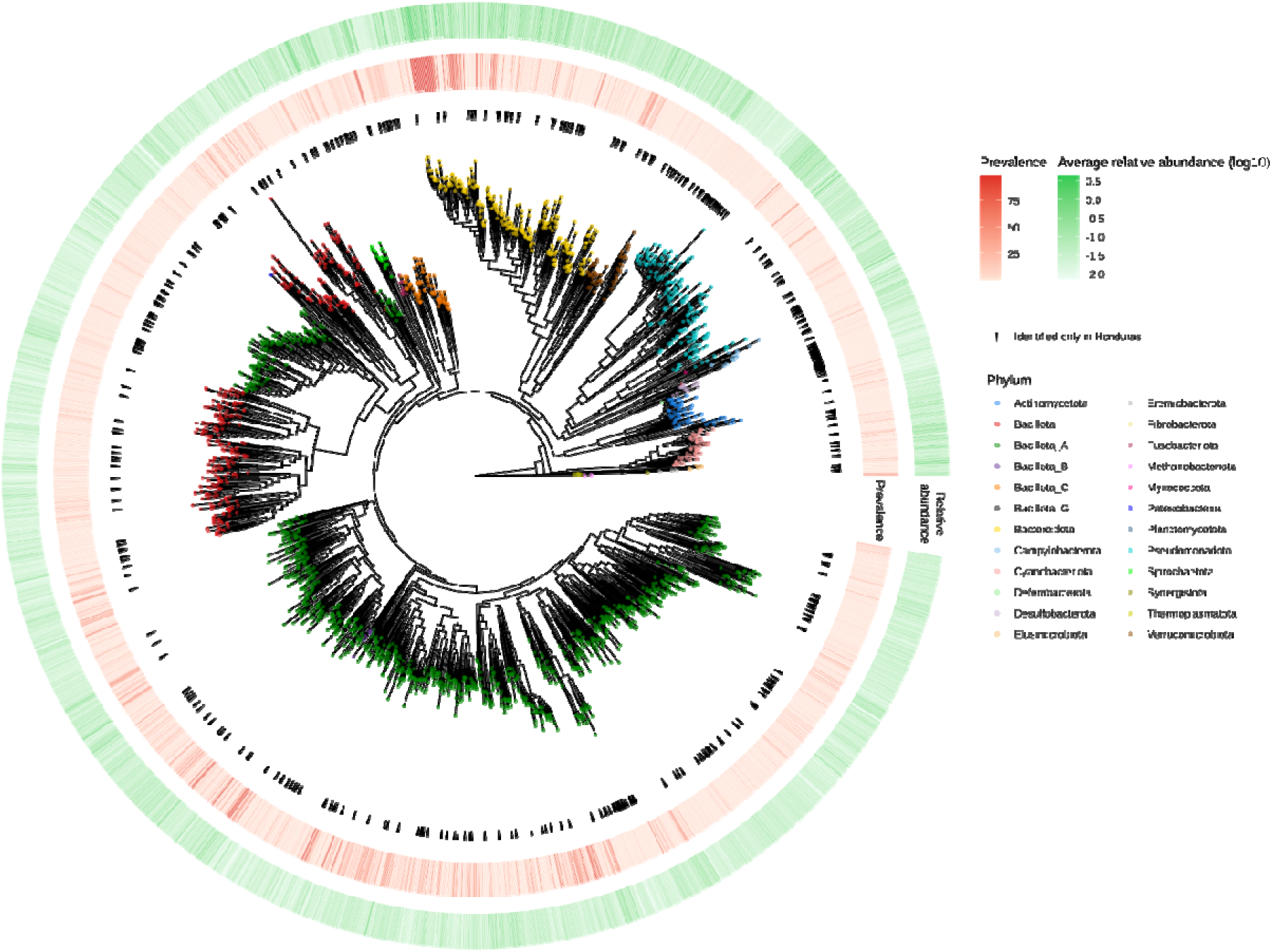
Phylogenetic tree of the 2,609 reconstructed MAGs representatives. The tree was built using PhyloPhlAn 3 using the PhyloPhlAn markers. Tips are annotated according to phylum level taxonomy predicted by GTDB-tk, and relative abundance, prevalence in the Honduras dataset are reported in the two outmost ring annotations. Species identified exclusively in this dataset are marked with a triangle.

We then further improved the characterization of our cohort by employing metagenomic assembly in order to reconstruct metagenome-assembled genomes (MAG). Samples were assembled with metaSPADES ^45^, binned with VAMB^46^, and quality checked with CheckM2^47^ (see Methods). After quality control, we recovered a total of 130,852 MAGs meeting the medium-quality thresholds proposed by MIMAG (>50% completeness and <10% contamination), with 63,459 (48.5%) of them being classified as high-quality (>90% completeness and <5% contamination) ^48^ (Figure S6, Table S4). After deduplication, this yielded a total of 2,609 representative genomes.

The species with the greatest number of genomes reconstructed is *Agathobacter faecis*, previously known as *Roseburia faecis*, with 1,494 genomes reconstructed and a prevalence of 91%. *Prevotella* was the genus with the largest number of reconstructed genomes (n species = 72, n genomes = 17,594), with several *Prevotella* species identified as the most prevalent in our dataset (Table S5, Figure S7). Excluding species from the *Prevotella* genus, the second most prevalent species was identified as *Faecalibacterium longum*, with a prevalence of 93%. Genus CAG-269 from Clostridia showed the largest number of species-clusters (n species = 74, n genomes = 1,641). Bacterial genomes spanned 22 different Phyla, with most of the species assigned to Bacillota A (1365), followed by Pseudomonadota (207), Bacteroidota (271), and Bacillota (380).

Archaeal organisms were reconstructed as well, for a total of 9 species from the Thermoplasmatota (6), Methanobacteriota (2), and Thermoproteota (1) Phyla. The most prevalent archaeal organism was identified in *Methanobrevibacter smithii*, which was found in the 51% of the samples and had 803 reconstructed genomes. We identified a second *M. smithii* clade as well, classified by GTDB-tk as *Methanobrevibacter smithii* A, and later proposed to be renamed as *Candidatus Methanobrevibacter intestini* ^49^. Such species were identified in nearly 30% of the samples, and we observed (in 716 samples) a mutual exclusion between *M. smithii* and *C. M. intestinii*.

To explore the degree of novelty of our reconstructed MAGs, we used two approaches to estimate how many species were not reported in other databases. First, we evaluated the taxonomy assignment from GTDB-tk^50^, and then we quantified the overlap between our reconstructed set of genomes and 289,232 MAGs from the Unified Human Gastrointestinal Genome (UHGG) ^51^ catalog. We identified 861 species that lacked a representative from UHGG results to date (Figure S8), being only observed in our cohort. GTDB-tk failed to assign species-level taxonomy for 1,976 genomes, revealing 391 unknown species spanning 3 families and 19 genera. Most of those species were retrieved from class Clostridia; in particular, we reconstructed 215 genomes from a novel species classified under the RUG11200 genus.

Among the reconstructed species in the Clostridia class, we were able to reconstruct 109 genomes for SGB 41716, the species identified by MetaPhlAn as the most prevalent. SGB4716, identified by MetaPhlAn as *Lacrimispora amygdalina,* was annotated by GTDB-tk as uncharacterized *Alitiscatomonas* sp900066535, with 77.56 % average nucleotide identity with *Lacrimispora amygdalina*. Metabolic pathways reconstruction and metabolite production prediction showed that the species is a producer of short-chain fatty acids (SCFAs), with model-based evidence of production of butyrate and propionate. We analyzed the population structure of the species by performing a pangenome analysis of the reconstructed genomes. Principal Component Analysis (PCA) highlighted the difference in functional potential of the strains of *Alitiscatomonas sp.,* with strains retrieved from Honduras separated in the PCA space (Figure S9). Metabolic features are distinguishable between strains, with speciating gene family variants retrieved from Honduras and other non-Western datasets involved in the pentose-phosphate pathway (K07404) and the TCA cycle (K01681), as well as a LysR binding domain and a dicarboxylate carrier protein (Figure S10).

In addition, we performed gene prediction and annotation on the reconstructed contigs. We built a gene catalog starting from 244,932,121 predicted protein sequences that were clustered at 95% amino-acid identity, resulting in 9,883,899 gene families. Among those, 413,626 gene families (4.2%) were annotated with a COG category, although the majority were of unknown function (S category, n= 1,514,506) or had no hit at all (n= 3,470,273). This fraction could represent protein fragments and proteins of non-bacterial origin (e.g., viral, eukaryotic), although 25,535 proteins of eukaryotic origin were detected, mostly from micro-eukaryotes and food.

When combined with gene families reported in the UHGG, we observed that 3,508,259 gene families have no representation within the UHGG protein database. Functional annotation of such gene families with eggNOG showed that 171,380 gene families (4.8%) lacked functional annotation, as extrapolated from COG functions and PFAM annotations. We further investigated the functional nature of those gene families and identified gene families that lacked any type of functional annotation, including a broader PFAM annotations. For those with some level of annotation, the majority of the proteins were annotated as part of COG categories M (Cell wall/membrane/envelope biogenesis), L (Replication, recombination and repair), and K (Transcription).

### Non-bacterial components

We explored the composition of non-bacterial components of the microbiome. To test for the presence of eukaryotic microorganisms, stool samples were screened for presence of ova and parasites. The exam showed the colonization by several organisms such as *Ascaris lumbricoides* (14%, 167/1148), *Entamoeba coli* (14%, 165/1148), *Entamoeba histolytica* (13%, 155/1148), *Blastocystis spp.* (5%, 67/1148), *Trichuris* (2%, 28/1148), and *Iodamoeba bütschlii* (1.5%, 17/1148). Polyparasitism was observed in 151 individuals, mostly occurring with species *Entamoeba coli* and *Entamoeba histolytica*, followed by *Entamoeba coli* and *A. lumbricoides*. The presence of *Entamoeba coli* and *I. butschii* in the human gut has been previously reported as nonpathogenic and usually occurs at low prevalence. Prevalence of *Blastocystis spp.* is surprisingly low in our population compared to previous reports in Central America and other non-western countries ^52–55^. High colonization by soil-transmitted helminths like *A. lumbricoides* and *Trichiuris* is commonly observed in subtropical countries and is one of the greatest public health concerns in developing countries ^56^. Despite higher rates of helminth infections, very few participants reported taking antiparasitic medications prior to our evaluation (*n*=2).

By using the binned contigs, we screened for the presence of genomes of eukaryotic origin. Recovery of eukaryotic genomes was performed with EukRep ^57^, which identified a total of 129 bins (Table S6-7). Quality control with BUSCO identified a total of 55 bins assigned to the Stramenophiles lineage. Further taxonomic assignment was performed by MMseqs2^58^ which determined that all the 55 bins were classified in the *Blastocystis* genus, a prevalent inhabitant of the human gut ^53,59,60^. Using the 55 recovered new eukaryotic genomes, alongside a catalog of eukaryotic genomes, we proceeded to get estimates of relative abundances of those organisms (Figure 4C). A total of 13 organisms were identified by mapping the metagenomic reads against the genome catalog using coverM. The most prevalent organism detected was *Blastocystis spp.*, present with multiple subtypes (ST1 n=530, ST2 n=303, ST3 n=577, ST4 n=3, ST6 = 1, ST7 n=1, unknown subtype n=79), followed by *Giardia intestinalis* (n=17) and *Saccharomyces cerevisiae* (n=12). We were not able to find most of the organisms that were discovered though microscopy assessment of the fecal material when using metagenomic mapping.

**Figure 4:**
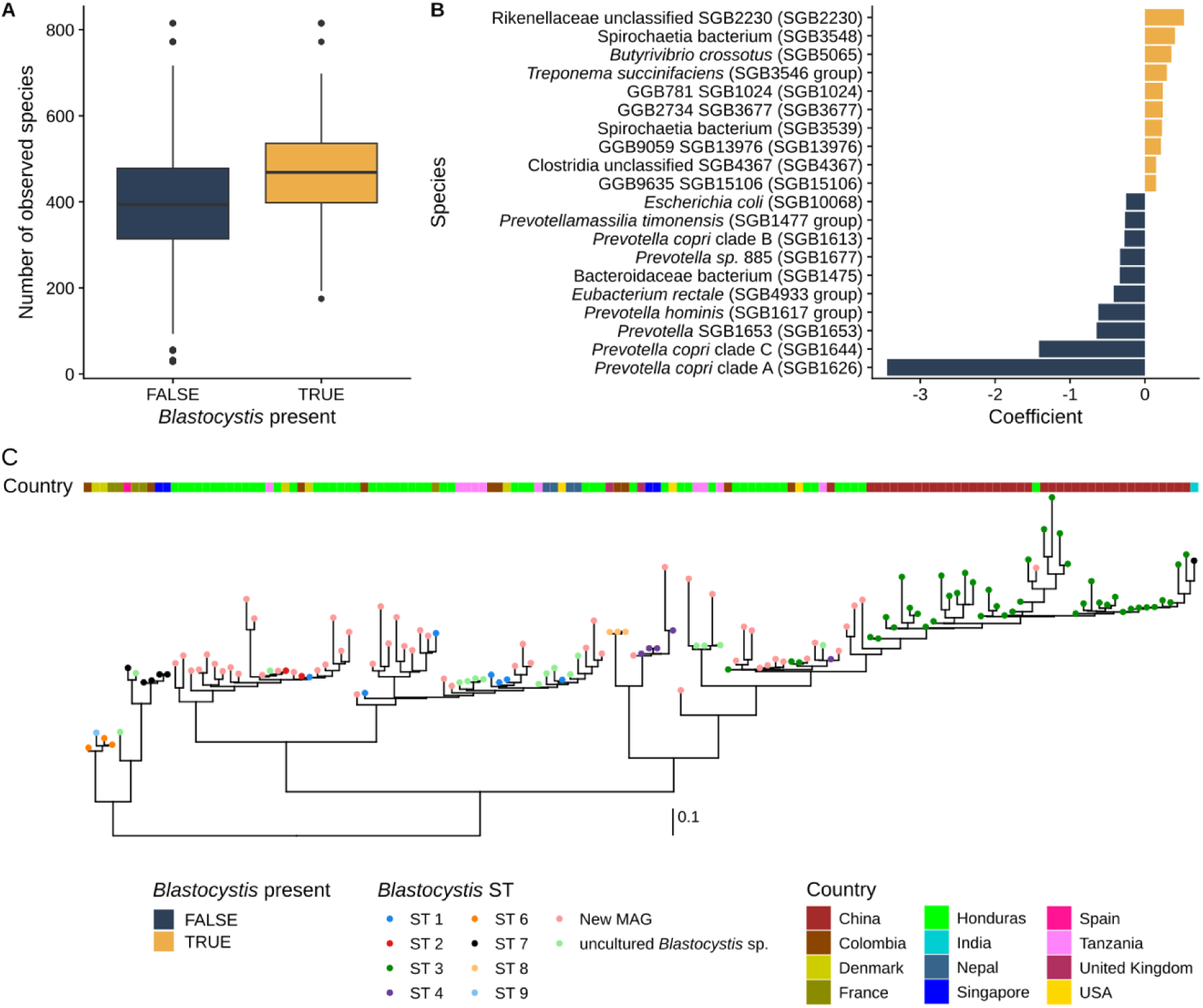
We identified *Blastocystis* in 1,010 individuals and observed that carriers have higher alpha diversity compared to non-carriers (A). Additional differences between carriers and non-carriers can be found in microbiome composition, as linear models identified species with differential abundances (B). We reconstructed 55 MAGs classified as *Blastocystis spp*. And built a phylogenetic tree of the new MAGs alongside publicy available isolate genomes (C). Subtype assignation was performed by ANI clustering with isolates retrieved from NCBI. The tree was built using PhyloPhlAn 3 using Stramenphiles BUSCO markers. MAGs retrieved by this study are labeled as ‘New MAG’.

Consistent with what has been previously reported, we observed a higher prevalence of individuals carrying at least a *Blastocystis spp.* subtype (*n* = 1,010), and 225 individuals were carriers of multiple subtypes. The overall cohort prevalence is 56% – higher than other countries in which *Blastocystis spp.* prevalence has been investigated ^53,60^. Per-village prevalence ranges from 45% to 70%, and the subtype distribution resembled the cohort distribution.

We examined the differences between the microbiome composition of *Blastocystis spp.* carriers and non-carriers and found that those who carried *Blastocystis spp.* had higher alpha diversity than non-carriers (Figure 4A, Wilcoxon rank-sum test observed species *P* = 1.38×10^−94^; Shannon diversity *P*=1.53×10^−87^). We expanded our analysis by looking at associations between *Blastocystis spp.* carriage and overall microbiome composition (Figure 4B, Table S8). The overall microbiome composition can be differentiated for carriers and non-carriers, with an increased abundance of taxa like *Butyrivibrio crossotus*, *Treponema succinifaciens,* and *Methanobrevibacter smithii* in carriers. In contrast to previous reports ^53,61^, we observed depletion of several *Prevotella* species and *Eubacterium rectale* in non-carriers. This reflects similar results previously observed in a cohort in Colombia ^62^. We then looked for associations with individual subtypes and bacterial species. Overall, we did not find any changes in the direction of association when looking at individual subtypes, with 28 species associated with one *Blastocystis spp.* subtype and 15 species associated with two or more subtypes (Mixed Effect Linear Model *q-value* < 0.05, *aabs ()* > 0.5).

Long-term colonization of *Blastocystis spp.* was observed in our 301 longitudinally sampled individuals, in which 71 individuals were found to carrying *Blastocystis spp.* after two years. Moreover, 5 individuals carrying multiple subtypes were still carrying the same subtypes of *Blastocystis spp.* after two years, with *Blastocystis spp.* ST3 being the most prevalent subtype among long-term carriers (n=41).

Next, to characterize the composition of the gut virome, all bins produced by VAMB were post-processed by the PHAMB pipeline, and, after quality control, we obtained 110,089 viral MAGs that were clustered into 29,958 viral operational taxonomic units (vOTUs) (Table S9). In addition to those MAGs, we obtained 12,136 viral clusters from contigs that were classified as a provirus by CheckV^63^ and that were clustered separately.

A consecutive clustering with representative sequences from the MGV database highlighted that the great majority of the sequences were of unknown origin, and only 3,353 newly reconstructed vOTUs clustered with a MGV reference sequence and 25,580 novel vOTUs with respect to MGV. For almost all vOTUs, geNomad^64^ was able to assign them to the *Caudoviricetes* class, with some of them annotated as families *Circoviridae*, *Crassvirales* and *Microviridae*.

By using Clustered Regularly Interspaced Short Palindromic Repeats (CRISPR) spacers, we investigated the associations between microbes and viral genomes by predicting CRISPR spacers in all our reconstructed MAGs. This resulted in 104,851 CRISPR spacers that further deduplicated into 4,935 spacers. Cluster representatives were mapped against all our viral MAGs, and 48,862 spacers mapped against a viral MAG enabling further host-association analysis for 2,417 vOTUs. A total of 403 vOTUs that associated with a bacterial host were associated exclusively with one bacterial species; 541 vOTUs were associated with more than 2 and less than 10 hosts; and 83 vOTUs with 10 or more hosts. We also observed the presence of vOTUs associated with different species across different family, class, phylum, and some occurrence of trans-kingdom association, as 5 associated with both bacterial and archaeal species. The majority of the vOTUs were associated with *Prevotella* species (*n*=326) as their host, followed by the *Alloprevotella (n=*37*), Coprococcus* (*n*=34), and *Clostridium* (*n*=33). Among the vOTUs that were represented in MGV, we observed a similar pattern in assigned taxonomies, with vOTUs with *Prevotella* species as the predicted host being the ones present in higher abundance and higher prevalence.

We mapped the vOTU representatives to the original metagenomes to produce relative abundance profiles. Overall, viral reads accounted, on average, for 4.44 % of the original metagenomes (1.68% sd); individuals showed a median value of 130 vOTUs present in each individual (mean 142.4, s.d. 66.13). Viral diversity was found to be positively correlated with bacterial diversity, as measured as number of identified species (Spearman correlation ρ = 0.73). We observed as well that many viral OTUs were private, i.e. present exclusively in the individual from which the viral MAG was originated.

A total of 255 vOTUs were annotated as Crassvirales, the family of crAss-phage, a gut microbiome component found to be globally widespread, and were found in 1,129 samples, roughly 50% of the total, with an average abundance of 5.5%. Using skani, we then aligned the representative sequences of the identified Crassvirales vOTUs to the crass-phage contigs identified and annotated from prior work ^65^. The crass-phage catalog contains 673 genomes, and almost half of the vOTUs identified in our dataset were represented by one of the already known crass-phage genome. Among these, we observed that most individuals have vOTUs assigned to the Candidate Family DeltacrAssvirinae (n=42), followed by Beta (n=32) and Epsilon (n=23).

**Figure 5:**
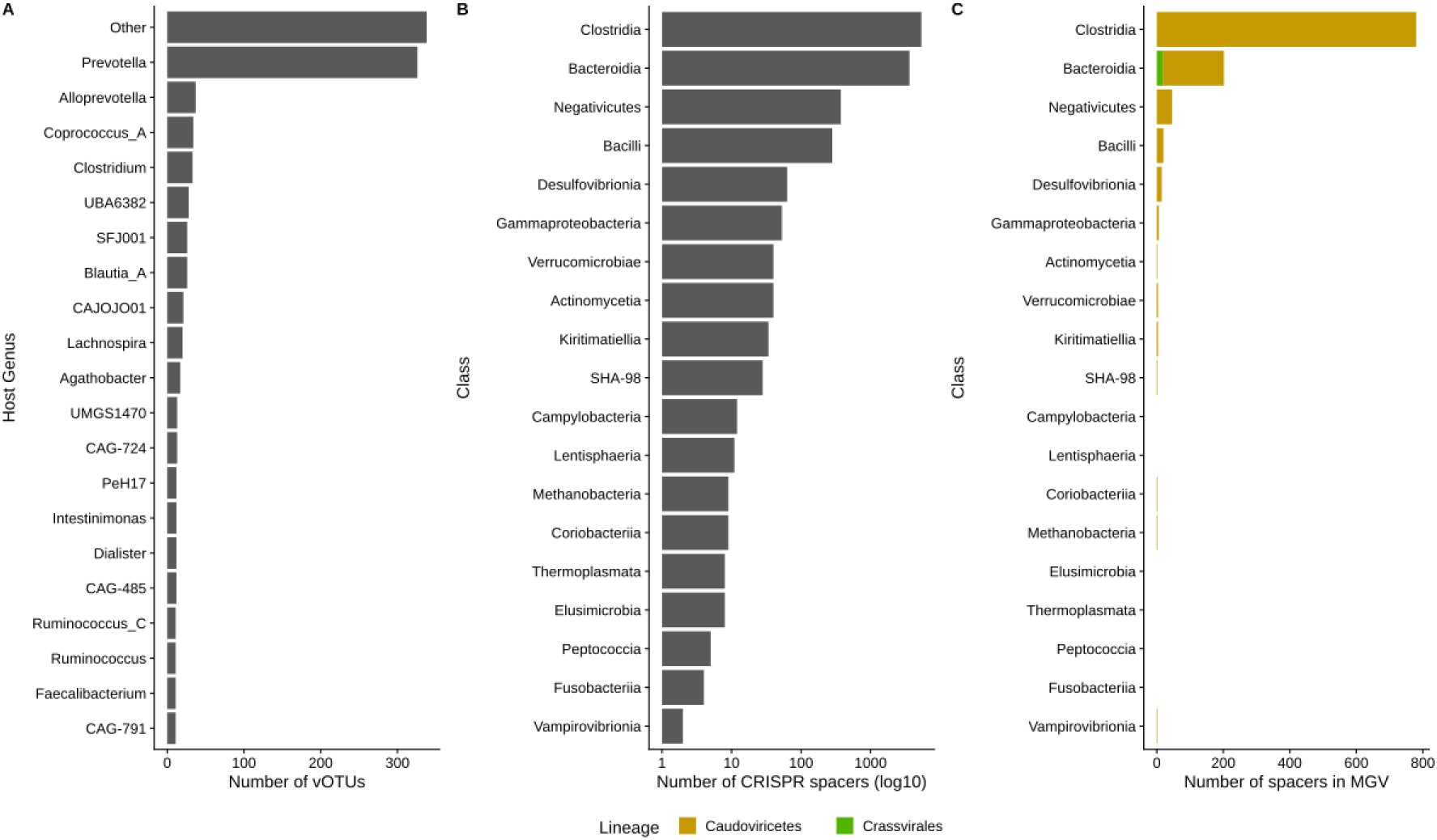
(A) Distribution of the predicted host genomes for the representative sequence of all the vOTUs. As the most prevalent bacterial genera, *Prevotella* is also the genus with most of the vOTUs are predicted to be hosted. Host prediction was performed by mapping the predicted CRISPR spacers to the sequence of the MAGs, we identified 48,862 CRISPR spacers, most of them were detected in *Clostridia* genomes (B), across all the genera, very few CRISPR spacers were identified in other public databases (C), with some genera without any representation.

### Longitudinal analysis of the gut microbiome

Prior work has documented the overall stability of the microbiome in the short and long term, with the ability to recover after exposure to microbiome-altering drugs, changes in diet, or colonization by pathogens ^66–68^. We investigated changes in the gut microbiome for a subset of individuals (n=301) who were re-sequenced after a period of two years. We compared the within-individual composition at both the taxonomic and functional level by assessing the stability of the microbiome using several approaches.

First, we investigated changes in composition and diversity, and we observed a significant decrease in alpha diversity measured with Shannon diversity and with the number of observed species between the two timepoints (Wilcoxon Rank-sum paired test, Shannon diversity p= 0.02535, number of species p=1.5e-07). We also found an overall change in composition structure (PERMANOVA p=0.007) assessed with a principal-coordinate analysis of the Bray-Curtis dissimilarity calculated on species-level relative abundances (Figure 6A). The variation of the microbiome structure across timepoints within subjects was less than that observed between individuals (within-subject versus between-subject Bray-Curtis dissimilarity Wilcoxon test: *P* <2.2×10^−16^, Figure 6C). We did not observe a significant change in the beta diversity of the functional potential, which was observed to be more stable than the taxonomic composition, suggesting preservation of microbiome functionalities.

**Figure 6:**
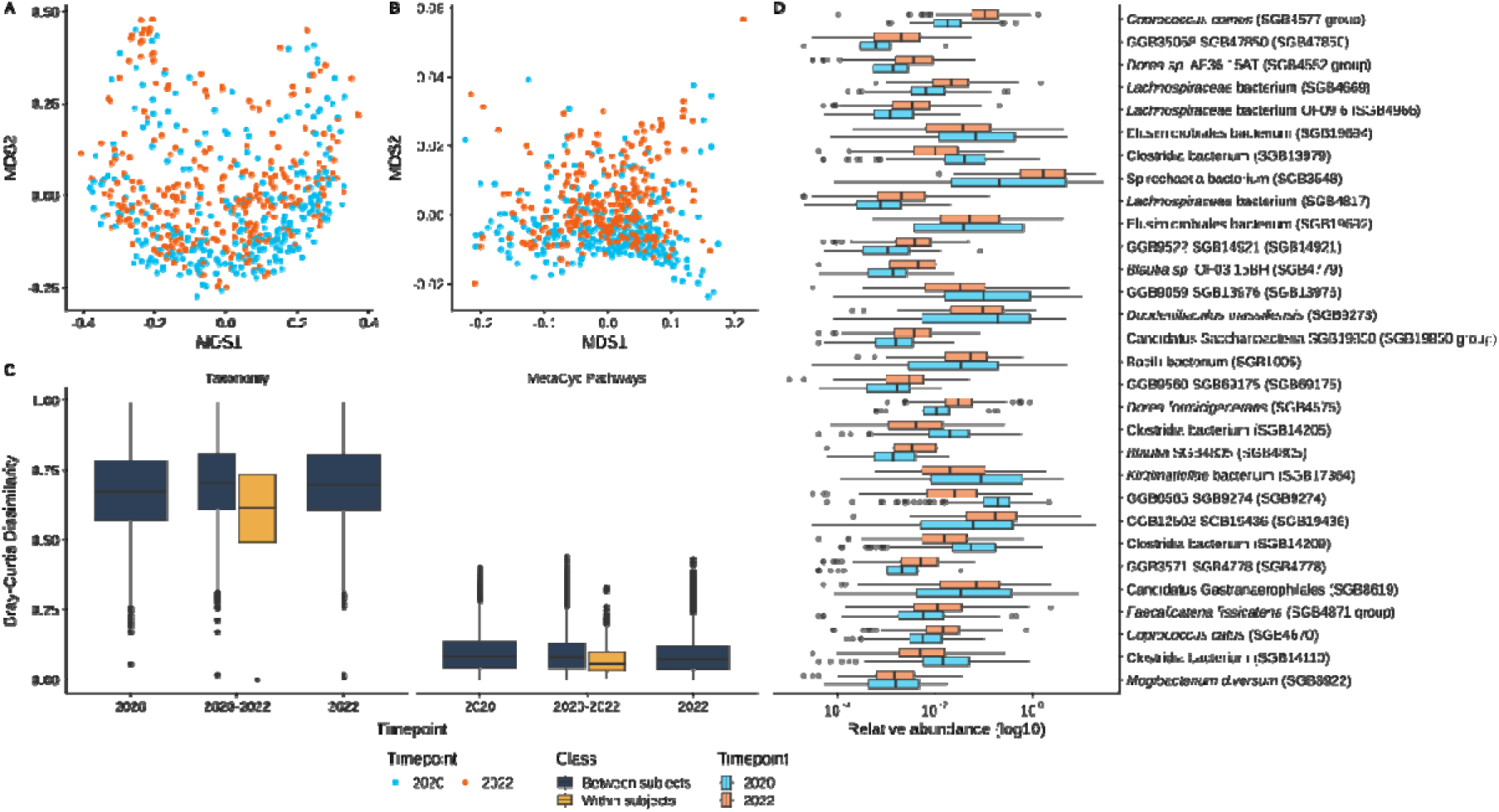
Principal Coordinate Analysis on the Bray Curtis dissimilarity calculated on species-level relative abundances (A) and MetaCyc Pathways relative abundance (B) show temporal separation between microbiome samples. Individuals were found to be more similar to each other after two years, but intra-individual beta diversity decreased (C). Linear mixed models as implemented by MaAsLin 2 identified 286 species differential abundant between individuals between and after infection with SARS-Cov-2.

Intra-individual bacterial species stability, measured with intra-class correlation coefficient (ICC) on most prevalent species (prevalence greater than 0.5) showed very low stability, with a median value of 0.09, with 17 species having an ICC value greater than 0.3. We then compared ICC values calculated on a longitudinal subset of samples from the HMP IBD dataset, including only samples from healthy donors,^16^ and showed higher species stability (ICC median 0.25, max 0.94).

We observed a similar pattern of changes in composition when looking at individual strain composition, observing a median strain-retention rate of 0.26 (lower than other cohorts ^44^). Looking into individual species retention rates, we found that several *Prevotella* strains are the most stable, with one out of three individuals retaining the same strain after two years.

This would suggest that the gut microbiome of a traditional-lifestyle non-western population might be more prone to variation over time, compared to western populations, given the fact that changes in medication usage, diet composition, or major diseases were not observed (although the intervening pandemic of COVID-19 may have played a role). Indeed, in recent years, there is growing evidence that infection with COVID-19 is associated with changes in composition in the intestinal tract. Individuals were asked to report whether they had an infection confirmed by a doctor; most of the individuals reported not having been infected with COVID-19 (n=296). We then compared the self-reported assessments with results from rapid immunoassay test to identify antibodies to SARS-Cov-2. A large fraction of the tested individuals (n=264) tested positive for the presence of IgG only, while only 3 individuals were positive for both IgG and IgM. Hence, we compared microbial profiles between the two timepoints using mixed-effect linear models as implemented in Maaslin2 (see Methods) to identify changes in species abundances and potential functional profiles. After controlling for individual ID as a random effect, and age, gender, and BMI as fixed effects, Maaslin2 identified 286 species differentially abundant between individuals before and after infection with SARS-Cov-2 (Figure 6D, Figure S11, Table S10-S11).

Among the statistically significant differential abundant species, *Coprococcus comes* and *Dorea* sp. AF36-15AT were the two species more enriched at the second timepoint among individuals positive for the presence of IgG only, while several *Spirochaetia* and *Elusimicrobiales* species were depleted among the same group of individuals. We also observed a major reduction in relative abundance of several species from the Prevotellaceae family, although they remained the most dominant taxa.

Finally, viral diversity was observed to be increased over the two-year period (Median T1 116 vOTUs, Median T2 131 vOTUs Wilcoxon Test P= 0.0005)

### Human genetic variation and microbiome associations

The genetic background of Honduras has been described as a mix of mostly Spanish, African, and native American ancestries, primarily Mayan^69^. Copán, the region in which our cohort is located, was the center of the Mayan civilization, and most of the people still identify themselves as Maya Ch’orti’.

Using low pass sequencing and successive genotype imputation, we performed genomic characterization of the individuals enrolled in our cohort. Most of the individuals (N=1,889) took part in the microbiome portion of the study, and, alongside the stool sample collection, they donated a saliva sample as well. The average sequencing coverage for the samples was 4X. The final dataset resulted in 29,693,799 SNVs identified in 1,701 samples, after retaining individuals with both human genome sequencing and shotgun metagenomic microbiome sequencing samples.

After filtering for minor allele frequencies, Hardy-Weinberg equilibrium, and performing LD pruning, we retained 227,216 SNVs, and we proceeded to compare human genetic variation with microbial variation. The first 10 genetic components from the PCA analysis explained most of the genetic variance in the dataset, and we proceeded to calculate the microbial variation that could be explained by host genetic variation in SGB and MetaCyc pathways composition. To reduce the total number of multiple tests performed, we filtered for SGB present in at least 20% of the population, retaining a total of 557 species. Using linear models, we fit each microbial species relative abundances, predicting it with the first 10 genetic components in order to calculate the percentage of explained variance (Figure S12, Table S12).

The total bacterial species variation explained by the genetic component was 2%, with 125 out of 557 species reaching statistical significance. One species in particular, SGB 4777 from the Lachnospiraceae family, reached an R^2^ of 9.7% (p-value from permutation test p=0.000001) and was found to be associated with the PC1, PC2, and PC8.

We performed the same analysis on 571 MetaCyc pathway abundances, which showed a smaller amount of total variance explained (0.5%), but a larger number of pathways reaching statistical significance (FDR < 0.05, n=289). The top pathways with higher R^2^ values are bacterial pathways involved in the biosynthesis of lipids, amino-acids, cofactors, and carbohydrates (Table S13).

We then examined the association between genetic variation and overall microbial diversity, as summarized by the Weighted UniFrac distance. The microbiomeGWAS tool^70^ identified 15 SNVs with MAF greater than 0.05 associated with differences in beta diversity (Table S14). Variants were identified close or within the genes DAB1, AP1S3, HTT, COX7B2, GABRA4, ARSB, RASGRF2, CALN1, NCKAP1L, DGKH, CYB5B (Figure S13, Supplementary Figure 9).

Next, to study the association between individual species and host variation, we performed a genome-wide association study on SNV with MAF greater than 0.05 and genus-level, species-level, and MetaCyc pathways abundances. We used linear regression as implemented in the MatrixQTL R package, including gender, age, BMI, and the first 10 genetic principal components as covariates (Figure S14, see Methods).

After combining genus- and species-level relative abundances and treating them as microbial traits, we tested 9,399,332 eQTL and identified 280 associations with FDR < 0.05. Only 7 eQTL reached the study-wide statistical significance threshold (p = 2.93 × 10^−11^) (Figure S15). Several SGB from the Lachnospiraceae and Ruminococcaceae were found associated with distinct SNV as well as the novel Ruminococcaceae genus GGB9729.

The strongest signal was identified on chr22:22208496, located within the IGL immunoglobulin lambda locus gene. Abundances of Lachnospiraceae SGB4817 were found to be decreased in individuals with one or two copies of the T allele.

When looking at the SNV identified as significant by microbiomeGWAS and by the eQTL model, we identified a total of 2 SNV in common between the two models, rs7679715 and rs17599102, both located on chromosome 4; rs17599102 is located on the GABRA4 gene and rs7679715 on the COX7B2 gene but still proximal to GABRA4. Individuals homozygous for both the rs17599102 A/A and rs7679715 C/C genotypes showed decreased median abundances of Ruminococcaceae SGB15234 and Clostridia SGB4372, and smaller beta-diversity.

We tested associations between SNV and pathways as well, with a total of 7,378,167 eQTLs. Although no eQTL reached the study-wide significance threshold, 119 were statistically significant with FDR < 0.05 (Table S16).

From metadata collected during the sample collection, we know about family members who are directly related to each participant, if at least one participant provided the information. Genetic sequencing allows us to further infer the degree of relationships of all participants in the cohort. Using KING, we calculated the pairwise kinship coefficient for each pair of participants (N= 1,622,078 pairs) and kinship values were grouped in degree of relation by using pre-calculated thresholds (See Methods). Participants in our cohort are, on average, related to each other at the second degree (□ = 0.1313), and this was observed when looking at the cohort as a whole and each separate village. Although some villages have both higher and lower kinship coefficients, we did not find any correlation with village size.

Using the weighted UniFrac distance and the strain-sharing rate calculated for all the pairs of individuals, we looked at the similarity of microbiome of people who are more genetically similar (Figure 7). The distribution of the microbiome similarity among pairs of people varies according to the three levels of relationship degree, namely (First to Second degree, Second to Third degree, and Third degree to unrelated), such that more genetically similar people have higher UniFrac distances (regardless of whether such a pair of people actually has a relationship) (Kruskal Wallis test P < 2.2 × 10^−16^). However, this analysis does not hold if we restrict the pairs of people to be village co-residents; in that case, perhaps because of the overwhelming shared environment, increased genetic similarity within a village is not associated with increased microbiome similarity.

**Figure 7:**
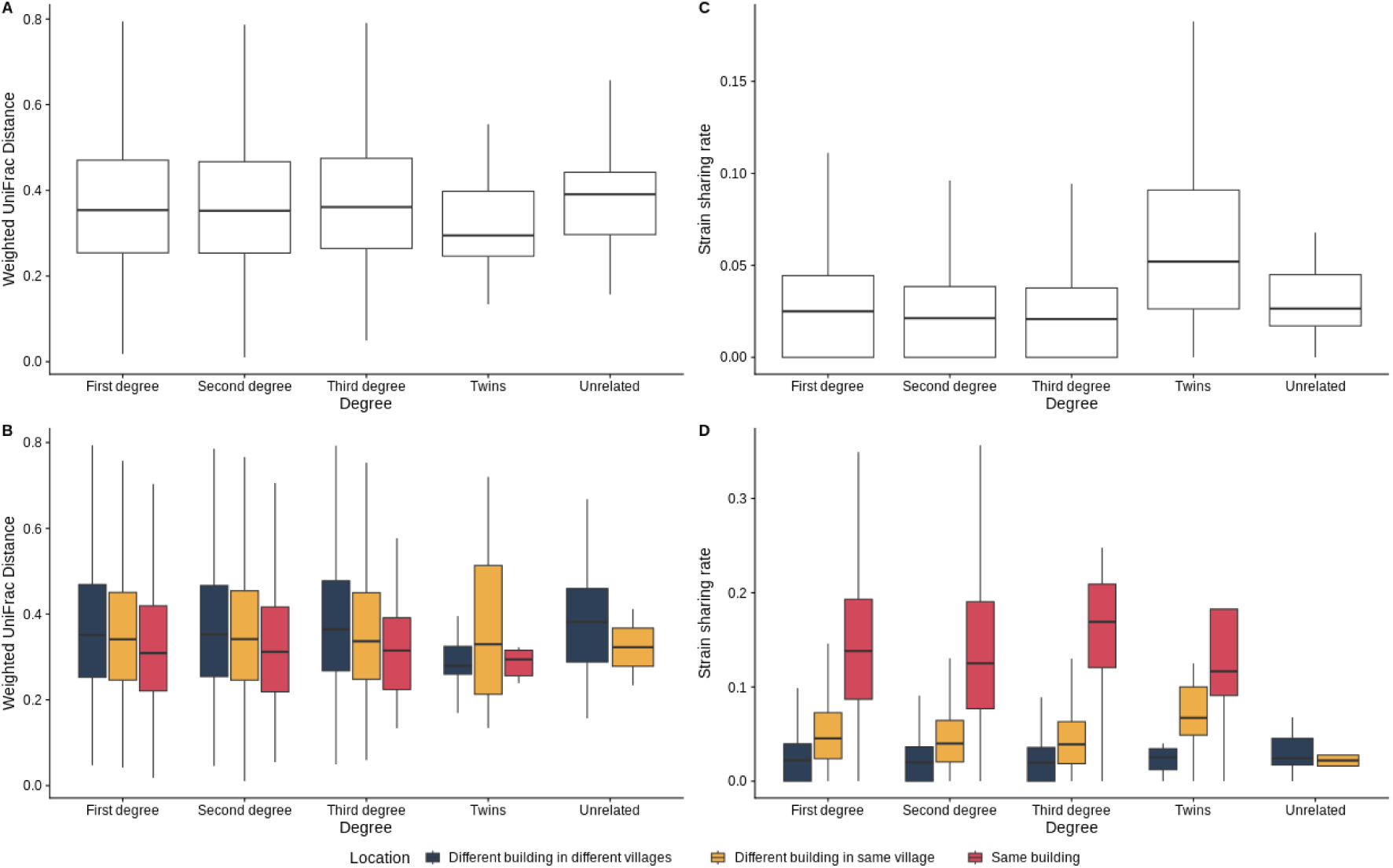
Distribution of the Weighted UniFrac Distance (A-B) and strain sharing rate (C-D) between different relationship degrees.

Although we have a very small set of individuals with a twin (N=45), we observed twin pairs having a more similar microbiome when they are in a different village than pairs living in the same village. In all the cases observed, the microbiomes of related pairs resemble unrelated individuals less (albeit with P=0.59).

We observed a similar trend when looking at the number of strains shared between individuals, namely the strain-sharing rate. Regardless their relationship degree, the number of strains shared across individuals living in different villages is relatively low, but it is higher in individuals living in the same village and in the same building. We observed an increased strain-sharing rate among third-degree-related individuals living in the same building, compared to first- and second-degree pairs. We hypothesized this was due to the inclusion of pairs of partnered individuals, as we previously observed higher relatedness between participants, but the median kinship coefficients among partners (□ = 0.1425) is closer to a second-degree relationship than a third-degree relationship.

When looking at actual relationships between individuals known to be socially connected (since we have detailed social network maps of the villages), and, after correcting for several covariates (such as BMI, age, gender, diet score), we observed that genetically similar individuals with an existing social relationship (such as a friend) have increased strain-sharing rate (Linear Mixed Model Kinship β = 0.11, *P <* 2 × 10^−16^; Relationship β = 0.84, *P <* 2 × 10^−16^; interaction between kinship and relationship β = 4.713 × 10^−2^, *P* = 1.42 × 10^−5^). In other words, information about genetic similarity of socially connected people further enhances the ability to predict their microbiome similarity, and this also holds even after accounting for diet, medication usage, socio-economic and demographic status (which have an effect on the existence of a relationship and not on the increase of shared strains).

In sum, overall, we find that genetic similarity, quantified by PCA analysis or by kinship, is associated with microbiome similarity.

## Conclusion

We present a comprehensive characterization of the gut microbiome of isolated, rural Honduran populations, significantly expanding our understanding of microbial diversity in non-western settings. Through shotgun metagenomic sequencing, we generated one of the most extensive microbiome datasets in Central America, uncovering novel bacterial species by reconstructing metagenome-assembled genomes (MAGs). These findings expand previous observations that traditional-lifestyle communities harbor unique microbial ecosystems that are distinct from those of industrialized populations, with numerous species that have either been lost or diminished in Western microbiomes, reinforcing the idea that modernization, via dietary changes and antibiotic usage, has contributed to a substantial reshaping of the human gut microbiome.

By integrating our dataset with global microbiome cohorts, we identified taxonomic and functional signatures that distinguish the Honduran gut microbiome from both Western and non-Western populations. We also expanded the Unified Human Gastrointestinal Genome (UHGG) by reconstructing and identifying novel prokaryotic species and proteins, and we showed that certain clades are exclusively present in our dataset.

In addition to prokaryotic diversity, we also explored the broader microbial ecosystem by looking at the eukaryotic and viral components of the gut microbiome. For instance, we identified and reconstructed 55 genomes of several subtype of *Blastocystis sp.*, a common gut protist. Notably, our findings suggest a higher prevalence of *Blastocystis* in individuals with greater microbiome diversity, reinforcing its potential role as a marker of gut ecosystem richness. We also reconstructed several thousand novel viral genomes, expanding the viral mappability of our metagenomes by over half, including novel bacteriophages and members of the *Crassvirales* family, which are thought to play critical roles in microbiome stability and bacterial population control.

The longitudinal component of our study provided insights into microbiome stability and temporal dynamics. Over a two-year period, we observed significant shifts in microbial composition, with notable changes in taxonomic diversity and functional profiles. While the gut microbiome exhibited resilience, the overall microbial diversity declined over time, a pattern that differs from observations in Western cohorts. This could be attributed to several factors, including possibly the SARS-CoV-2 epidemic, which caused significant microbiome alterations.

Moreover, our study investigated links between human genetic variation and gut microbiome composition. By leveraging low-pass human genome sequencing, we identified several microbial species and metabolic pathways that exhibit significant associations with host genetic components. These findings suggest that specific genetic factors may influence microbial colonization and metabolic potential, reinforcing the concept of microbiome adaptation to the host. Our genome-wide association analysis further identified specific genetic loci linked to microbiome variation, shedding light on potential host determinants of gut microbial diversity.

Finally, our investigation into microbiome transmission patterns provided compelling evidence for microbiome similarity among related individuals and those within close social networks. We observed higher strain-sharing rates among individuals living in the same households and villages, supporting the notion that microbiome transmission is influenced by both genetic relatedness and shared environments.

Overall, this study represents an important step forward in uncovering the vast microbial diversity of under-represented human populations. By characterizing the Honduran gut microbiome in unprecedented detail, we provide insights into microbial ecology and host-microbiome interactions. Our findings underscore the importance of expanding microbiome research beyond industrialized nations to gain a more comprehensive and global perspective on human microbial diversity. Future studies can continue to explore the interplay between environment and microbiome composition, with an emphasis on how these factors contribute to health and disease. Furthermore, the novel microbial species and functional pathways identified in this study offer promising leads for future research into microbiome-based therapeutics and precision medicine approaches tailored to diverse human populations.

## Methods

### Enrollment of participants

Our study was conducted in the department of Copán, Honduras, in an area of over 200 square miles of mountainous terrain with an estimated total population of 92 000 people. Villages and participants selected for this study were previously enrolled in the Honduras Social Network Targeting RCT project ^71^. Of 176 study villages, 18 villages were randomly selected to meet the study’s target size of 2000. Villages were chosen to vary by wealth, population size, elevation, isolation index, and time to the main road.

### Sample collection

Participants are instructed on how to self-collect the fecal samples using in-person education and a leave-behind training manual and asked to return samples to the local team or to the designated community liaison within 1hr of producing them. Fecal samples are collected using sterile collection tubes with a scoop. Samples were then stored in liquid nitrogen at the collection site and then moved to a −80 C° in Copán Ruinas, Honduras. Samples were then shipped on dry ice to the USA and stored in −80 C° freezers.

### Library preparation and sequencing

DNA was extracted using the Chemagic Stool gDNA extraction kit and libraries were prepared using the KAPA Hyper Library Preparation kit. Shotgun metagenomic sequencing was performed on an Illumina NovaSeq 6000 platform at a targeted sequencing depth of 50Gbp. Samples not reaching the desired sequencing depth were resequenced on a separate run.

Raw metagenomic reads were deduplicated using prinseq lite (version 0.20.2, ^72^) with default parameters. The resulting reads were screened for human contamination (hg19) with BMTagger ^73^ and then quality filtered with trimmomatic ^74^ (version 0.36, parameters ILLUMINACLIP:nextera_truseq_adapters.fasta:2:30:10:8:true SLIDINGWINDOW:4:15 LEADING:3 TRAILING:3 MINLEN:50).

### Read-based taxonomic and potential functional profiling

Quantification of organisms’ relative abundance was performed using MetaPhlAn 4^75^, which internally mapped the metagenomes against a database of ~5.1M marker genes describing more than 26~ species-level genome bins (SGB).

Microbiome species richness was estimated using the Shannon entropy index and the total number of observed (relative abundance greater than zero). Multidimensional scaling analysis (vegan cmdscale function) was performed on Bray-Curtis dissimilarity (vegan vegdist function) calulcated on the relative abundances obtained by MetaPhlAn4. Strain-level profiling was performed for a subset of species present in at least 10% of samples using StrainPhlAn 4 ^75^ using default parameters.

Phylogenetic trees generated by StrainPhlAn were annotated with country of origin or world region as a trait, and phylogenetic signal was calculated by using the phylo.d function as implemented in the caper R package.

Functional potential analysis was performed using HUMAnN 3.0 ^76^. Gene family’s profiles were normalized using relative abundances and collapsed into MetaCyc pathways and KO identifiers. CAZy profiles were obtained by regrouping the HUMAnN gene family profiles using a custom mapping file.

Differential abundance analysis was performed using mixed-effect linear models as implemented in Maaslin2 ^77^. Basic covariates (age, gender, BMI) were included in all models as fixed effects as well as village as random effect. Longitudinal models included the individual ID as random effect. Multiple hypothesis testing was performed using the FDR procedure.

A total of 30 additional studies comprising 10,173 human gut metagenomes were downloaded by accessing curatedMetagenomicData ^78^ and processed with the same pipeline used to process our collected data. An additional dataset from South Africa^29^, not present in curatedMetagenomicData as to date, was downloaded and processed as well.

### Metagenomic assembly and contig binning

De-novo metagenomic assembly was performed on all samples using metaSPADES ^45^ (version 3.13.1, default parameters), and contigs longer than 1000bp were discarded. Metagenomic reads were mapped against the assembled contigs using minimap2 ^79^ (version 2.24, parameters: -ax sr -t 12 -N50). The resulting alignments were used as input for the VAMB pipeline for contig binning ^46^ (version 3.0.9, parameters: --minfasta 500000). The automatic binning procedure generated a total of 226,039 MAGs that were then subjected to quality control to evaluate completeness and contamination using CheckM2 ^80^ (version 1.0.2). We selected a total of 130,852 MAGs passing the thresholds for medium-quality genomes proposed by another study ^48^ being at least 50% complete and displaying less than 10% of contamination.

MAGs were dereplicated at 95% ANI using galah ^81^ and cluster representative, selected using the formula proposed by one study ^82^, were taxonomic annotated with GTDB-tk ^50^ obtaining a total of 2,609 metagenomic species clusters. Raw reads were then mapped against the catalog of representative clusters with bwa-mem and the resulting alignments were processed by CoverM ^83^ to obtain relative abundances of the species clusters (parameters: min-read-aligned-percent 70, min-read-percent-identity 90, min-covered-fraction 10).

MAGs were annotated with Bakta^84^ (version 4.1, default parameters) using the “Full” database, and predicted proteins from each MAG were functionally annotated using eggnog-mapper ^85^.

Reconstruction of metabolic models was performed by gapseq ^86^ using the “doall” pipeline on each MAG, then species-level pangenomes were generated by “gapseq pan”. Prediction of metabolic by-products was performed using the R package cobrar.

Proteins predicted by Bakta were iteratively clustered with MMseqs2 ^58^ using the cluster pipeline to build non-redundant catalogues first at 100% sequence identity and 80% sequence coverage and then at 95% sequence identity and 80% coverage.

Species-level representatives identified by galah were used as input to PhyloPhlAn to generate a maximum-likelihood phylogenetic tree (parameters -d phylophlan --diversity high --accurate)

### Eukaryotic contig binning and prevalence estimation

Scaffolds generated by metaSPADES were screened for presence and recovery of eukaryotic contigs with EukRep ^57^ using default parameters. Samples with non-zero sequences were mapped against the original metagenome with minimap2 and then binned with metabat2. Eukaryotic genome completeness was estimated using BUSCO (v5) ^87^ using parameters ‘--auto-lineage-euk -m genome’ and the ‘eukaryota_odb10’ lineage.

The retrieved eukaryotic MAGs were used to build a minimap2 index and mapped against the original metagenomes using CoverM^88^ to estimate their relative abundance using parameters “---min-covered-fraction 80 --min-read-percent-identity 95 --min-read-aligned-percent 80”.

### Viral contig binning and prevalence estimation

Viral OTUs (vOTUs) were obtained by parsing the previously processed VAMB bins with PHAMB ^89^ and then quality checked with CheckV (workflow ‘end_to_end’) ^63^.

CheckV result tables were parsed in order to identify viral contigs. We retained contigs annotated as complete, medium- and high-quality contigs with ≥75% average amino acid identity (AAI), >30% alignment fraction of the contig, and less than 15% difference in genome size from theclosest CheckV database hit. Low quality contigs with more than 10 genes, 10% of which annotated as host genes, and 40% annotated as viral genes, were included as well to account for potential novel uncharacterized viruses. Proviral contigs annotated at least as medium quality were included as well.

Viral and proviral contigs were clustered and dereplicated at 95% ANI across 85% of the shorter sequence using the scripts provided by CheckV to generate vOTUs. Taxonomic assignation was performed by Genomad ^64^ using the end-to-end workflow. Additional taxonomic assignation was performed by clustering the vOTU representative sequencing with the MGV genome database using the same approach described above.

The representative sequences for the vOTUs were indexed with BowTie2 and mapped against the original metagenomes using CoverM ^83^ to estimate their relative abundance using parameters “---min-covered-fraction 80 --min-read-percent-identity 95 --min-read-aligned-percent 80”.

vOTUs discovered in contigs of bacterial MAGs were host annotated according to the bacterial host. The host taxonomy of other viruses was derived using CRISPR spacers. CRISPR spacers were internally identified by Bakta using PILER-CR. The combined set of CRISPR spacers mined from MAGs were then mapped to all vOTUs using blastn (v2.13). Subsequently, spacer hits were processed such that only CRISPR-spacer matches with ≥90% sequence identity over ≥90% of spacer length and a maximum of 3 mismatches were retained. Spacer hits were used to calculate the taxonomy at the species level on the predicted host bacteria using a majority rule.

### Human genome sequencing

Participants were instructed how to produce at least 5 ml of saliva. Saliva samples were subjected to low-pass genome sequencing on the Illumina NovaSeq 6000 after performing DNA extraction. Reads were mapped against the human genome (version hg38) using FQ2BAM as implemented in the NVIDIA Clara Parabricks pipelines, and the resulting BAM files were processed with GLIMPSE ^90^ to perform low pass imputation against the 1000 Genomes reference panel ^91^. Resulting VCF files were merged into a single BCF file with bcftools ^92^ and processed with PLINK ^93^ using parameters “--hwe 1e-3 --maf 0.05 --indep-pairwise 50 10 0.1”. This resulted into 13,651,831 variants across 2,240 individuals. We then applied filters to retain samples with paired metagenome and performed LD pruning, which yielded 1,701 individuals and 227,216 variants. Principal Component Analysis was calculated with PLINK using the “--pca” parameter.

Kinship analysis was performed using KING, and the ranges used for inferring the degree of relation from kinship coefficients were taken from the original publication.

### Association analyses

Microbiome association analysis was performed by fitting multiple linear models as implemented in Matrix eQTL ^94^ to predict microbiome features (species-level and MetaCyc pathways abundances) after including gender, age, BMI, and the first 10 genetic principal components as covariates. We included species present in at least 10% of the participants. SNPs identified as statistically significant were associated with the closest gene (nearest function in SummarizedExperiment R package). microbiomeGWAS^70^ was used to perform a GWAS on the Weighted UniFrac distance matrix, including the same covariate used for the microbiome QTL analysis. A GWAS using the beta-diversity distance matrix is built on the idea that if a genetic variant affects the microbiome, individuals who share more alleles at a specific locus (for example, both having AA means they share two alleles, while one having AA and the other GG means they share none) will have lower beta-diversity distances.

## Acknowledgments

We thank the NOMIS Foundation for support of this initiative. Additional support from the Rothberg Catalyzer, Paul Graham, and the Pershing Square Foundation. The underlying cohort was supported by the Gates Foundation.

## Author contributions

FB and NAC conceived and designed the study; NAC supervised the project; FB and NAC contributed to the methodology design and analytic approach; FB and NAC collected the data; FB performed the analysis and interpreted the data; FB, IB, MG, and NAC wrote the manuscript.

## Data Availability

Metagenomic reads are available on the NCBI Sequence Read Archive database with accession PRJNA999635.

