## Supplemental Materials for "Characterization of gut microbiomes in rural Honduras reveals novel species and associations with human genetic variation"

Supplementary Figures

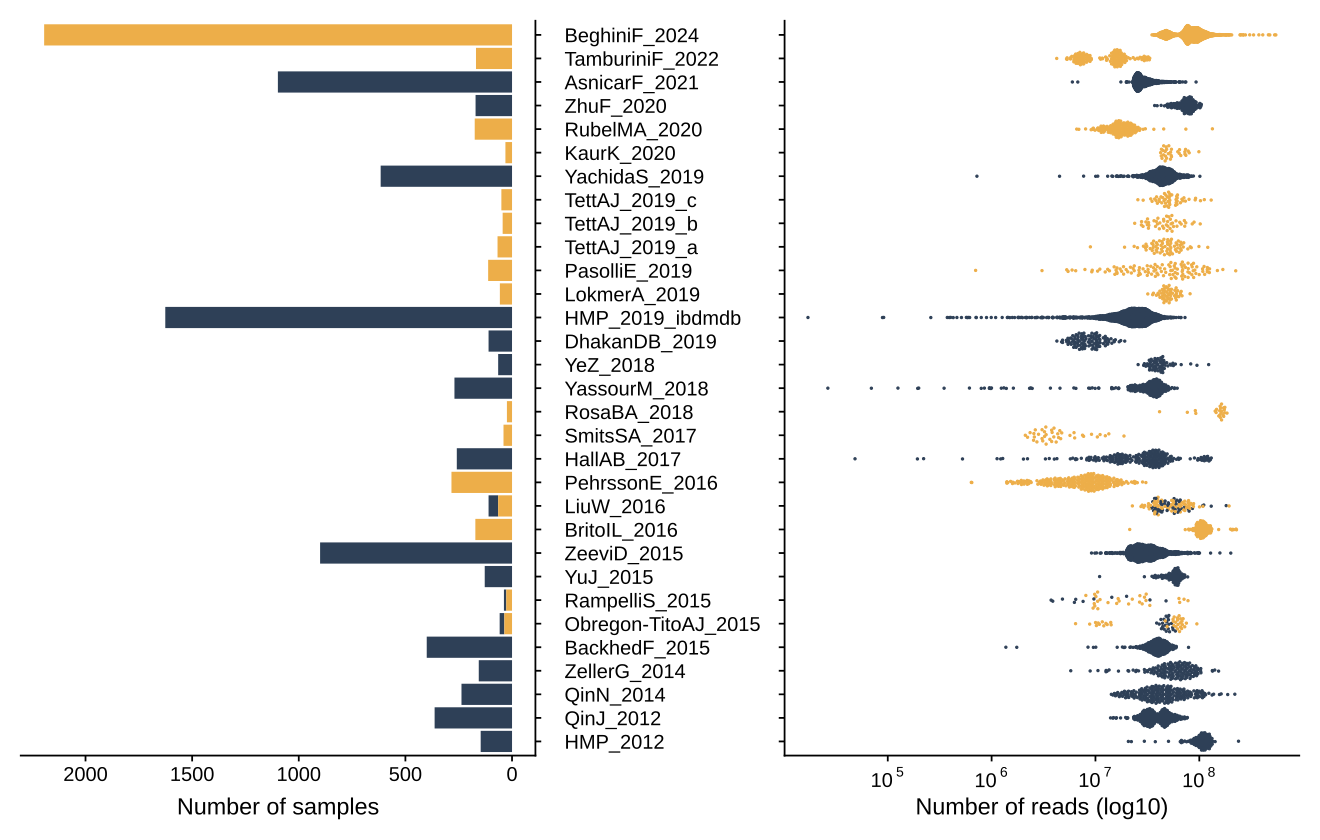

**Figure S1**: Characteristics of the 29 datasets and 10,173 samples considered in this study in terms of number of samples and metagenome size (as number of reads per sample). Bars and points are colored whether curatedMetagenomicData listed the sample and/or the dataset as westernized (blue) or non-westernized (yellow). The present dataset is in the top row.

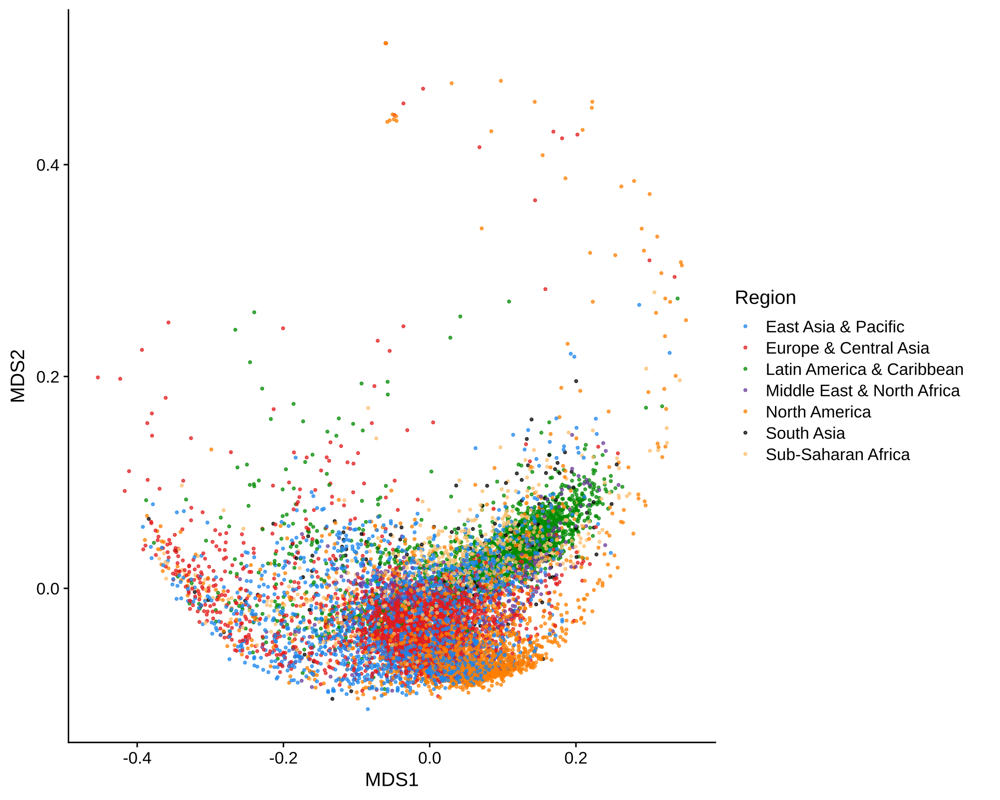

**Figure S2**: Principal Coordinate Analysis of the Bray-Curtis dissimilarity calculated on the MetaCyc Pathway’s relative abundance of 10,173 samples from curatedMetagenomicData and this study.

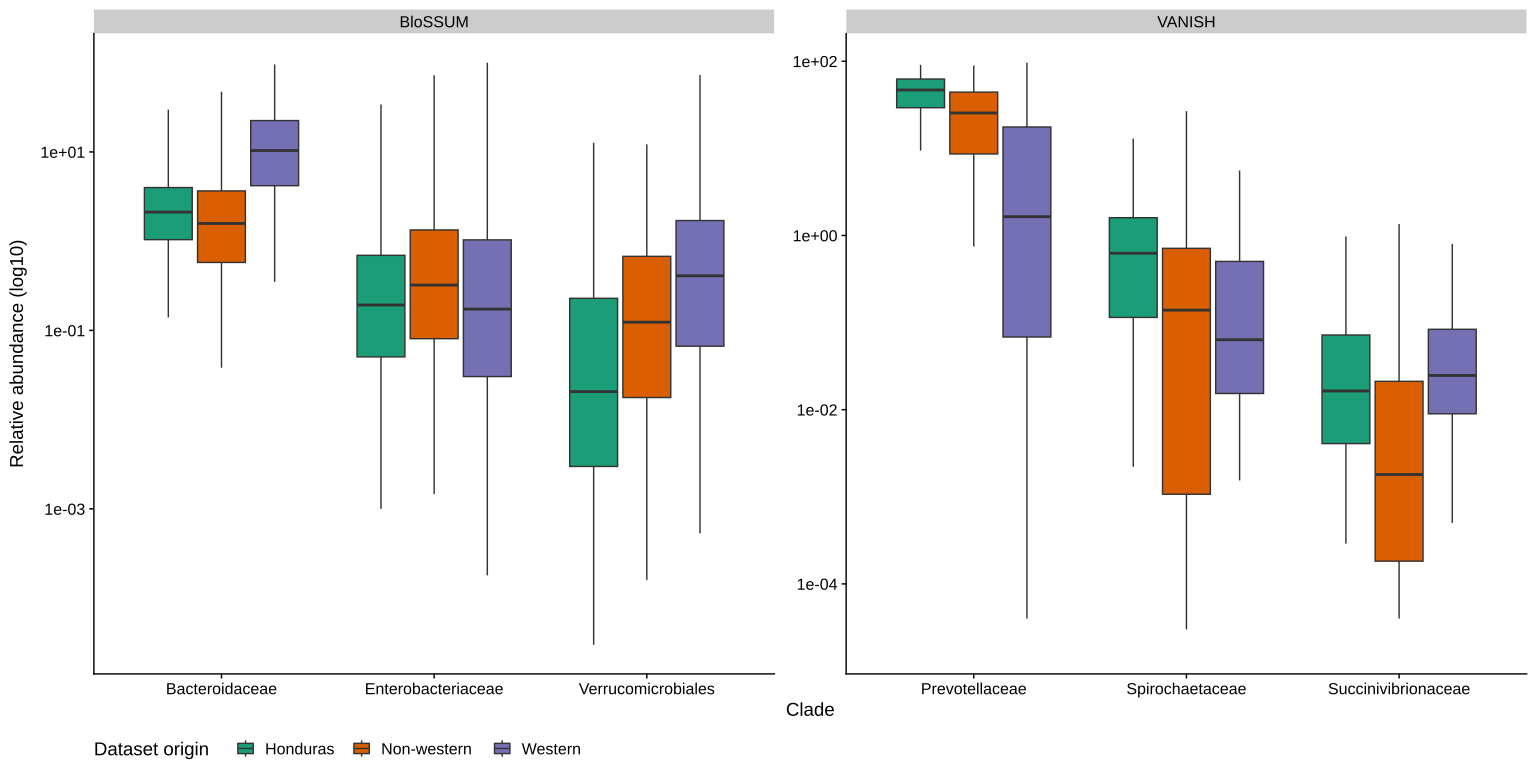

**Figure S3**: Distribution of relative abundance (log10) of BloSSUM and VANISH clades across dataset origins: Honduras (green), Non-western (orange), and Western (purple). BloSSUM taxa includes relative abundances of Bacteroidaceae, Enterobacteriaceae, and Verrucomicrobiales species, while VANISH taxa accounts for Prevotellaceae, Spirochaetaceae, and Succinivibrionaceae species.

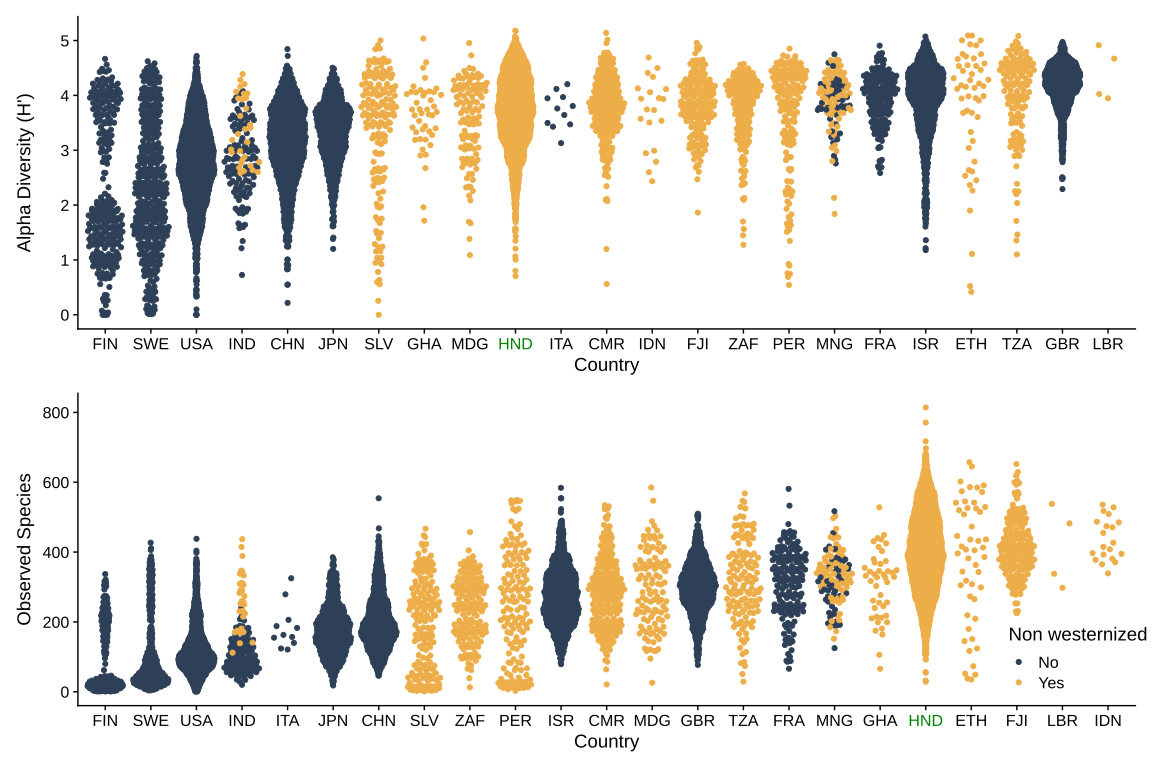

**Figure S4**: Distribution of the alpha diversity for 10,173 samples considered in this study calculated as Shannon Diversity (top) and number of observed species (bottom). Honduras (noted in the green color label) has more rich and diverse samples than other countries (P<0.05) except for Ethiopia, Indonesia, Cameroon, Fiji, Indonesia, and Liberia.

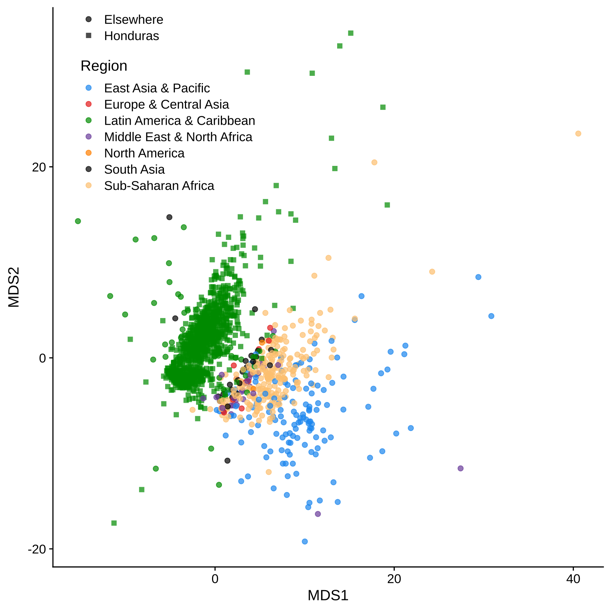

**Figure S5**: Principal Coordinates Analysis on the Kimura distance calculated on the phylogenetic tree reconstructed by StrainPhlAn for Prevotella copri clade H. Strains reconstructed from Honduras samples are spatially separated from strains reconstructed from other world regions (PERMANOVA P= 0.001).

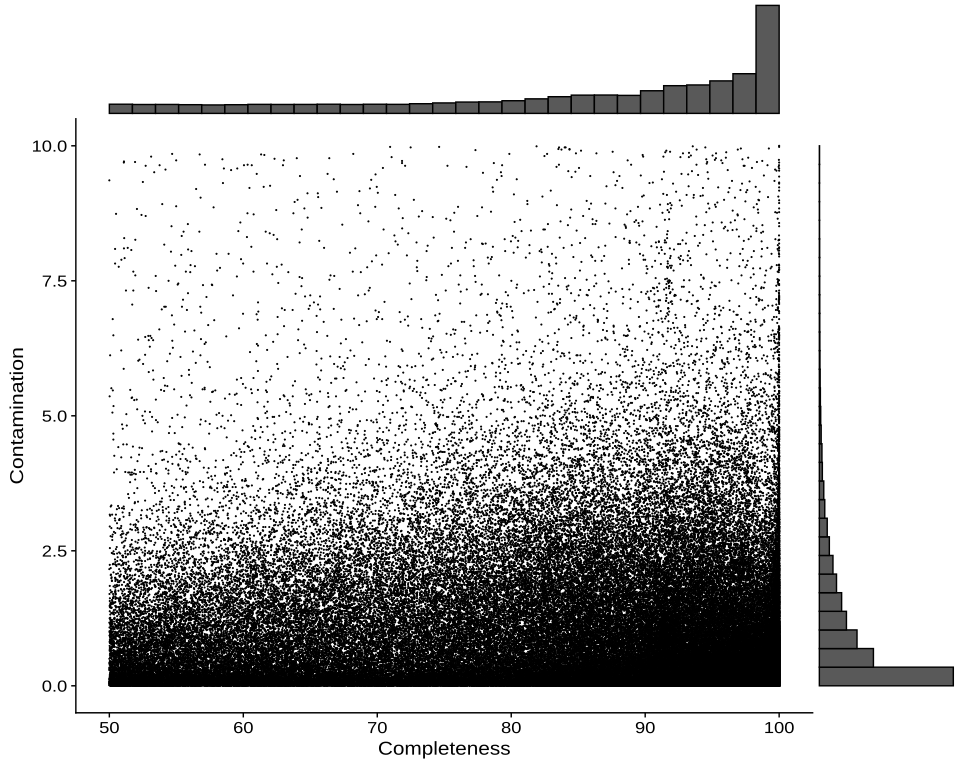

**Figure S6**: Distribution of completeness and contamination (as assessed by CheckM2) for all the 130,852 reconstructed MAGs meeting at least the medium-quality thresholds proposed by MIMAG (>50% completeness and <10% contamination).

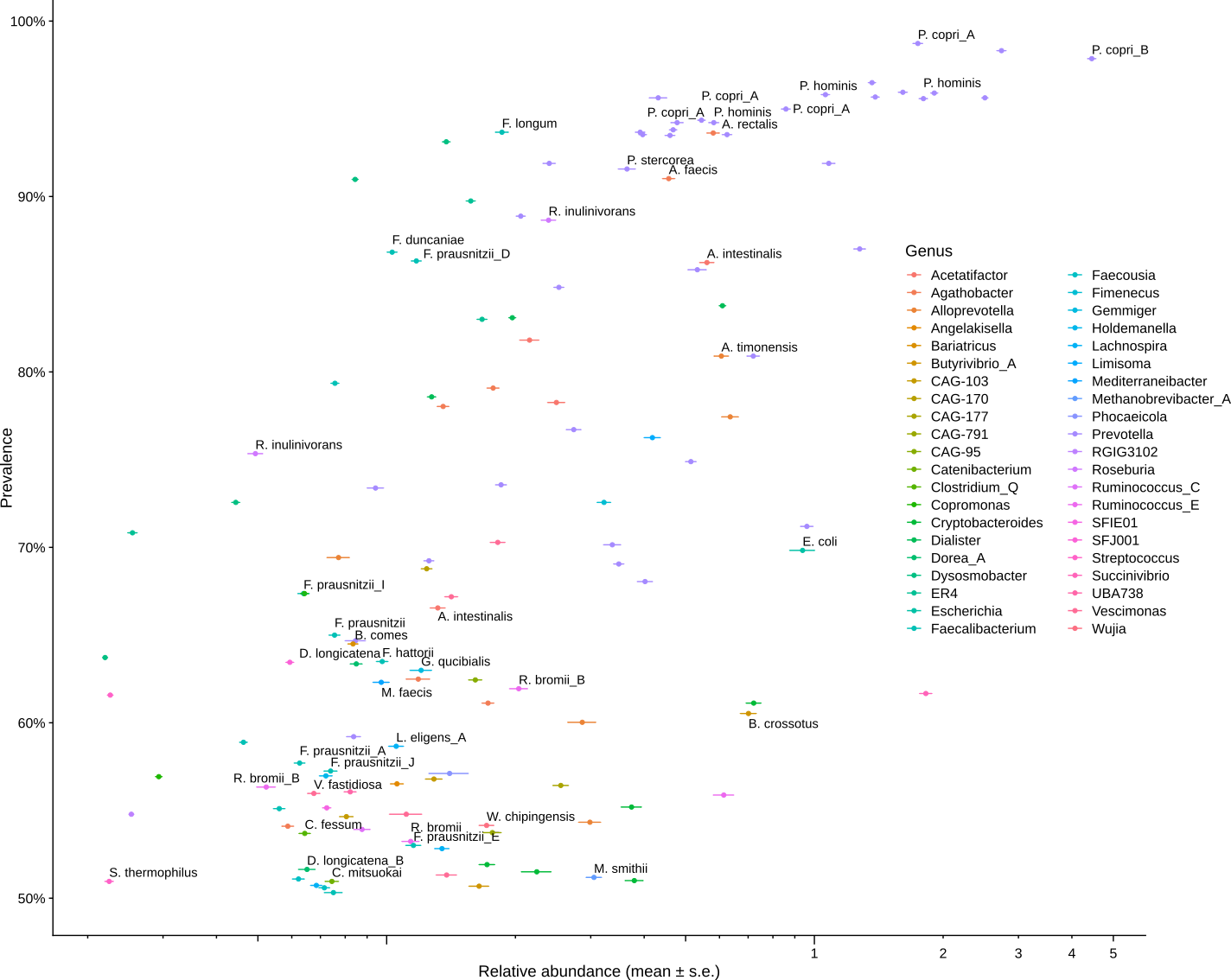

**Figure S7**: Scatter plot describing the relationship between species relative abundance and prevalence, as calculated from the CoverM relative abundances obtained by mapping cluster representative to the metagenomes.

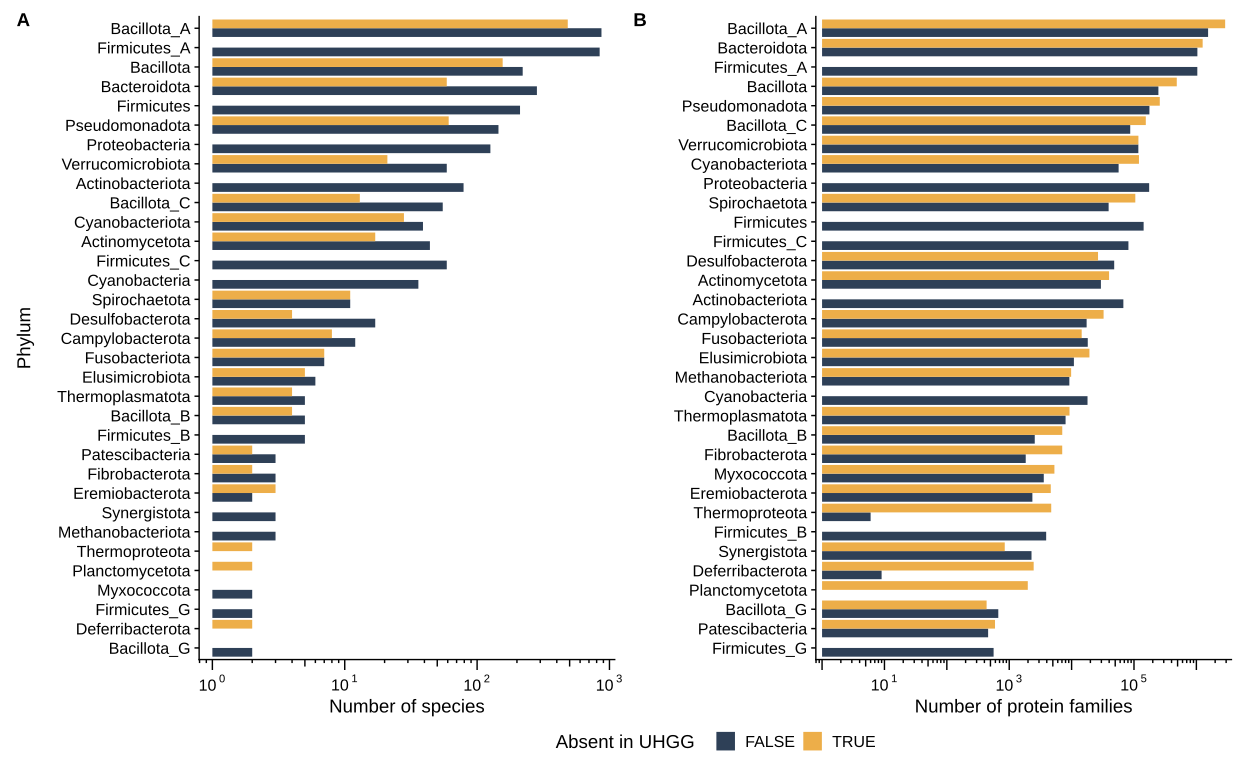

**Figure S8:** Distribution of species (A) and protein families (B) represented in the UHGG database. Presence in UHGG was determined by sequence clustering at 95% with UHGG genomes representative and non-redundant UHGG protein catalog. Protein families taxonomy was assigned by tracing back the taxonomy of the representative sequence.

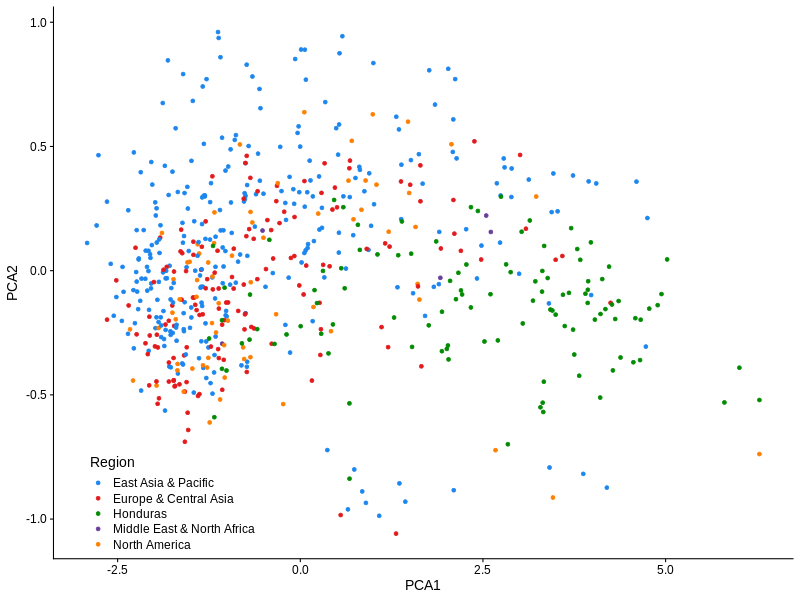

**Figure S9**: Principal Component Analysis on the Jaccard distance calculated on the presence/absence matrix of the pangenome of *Aliscatomonas* sp900066535 determined by panaroo.

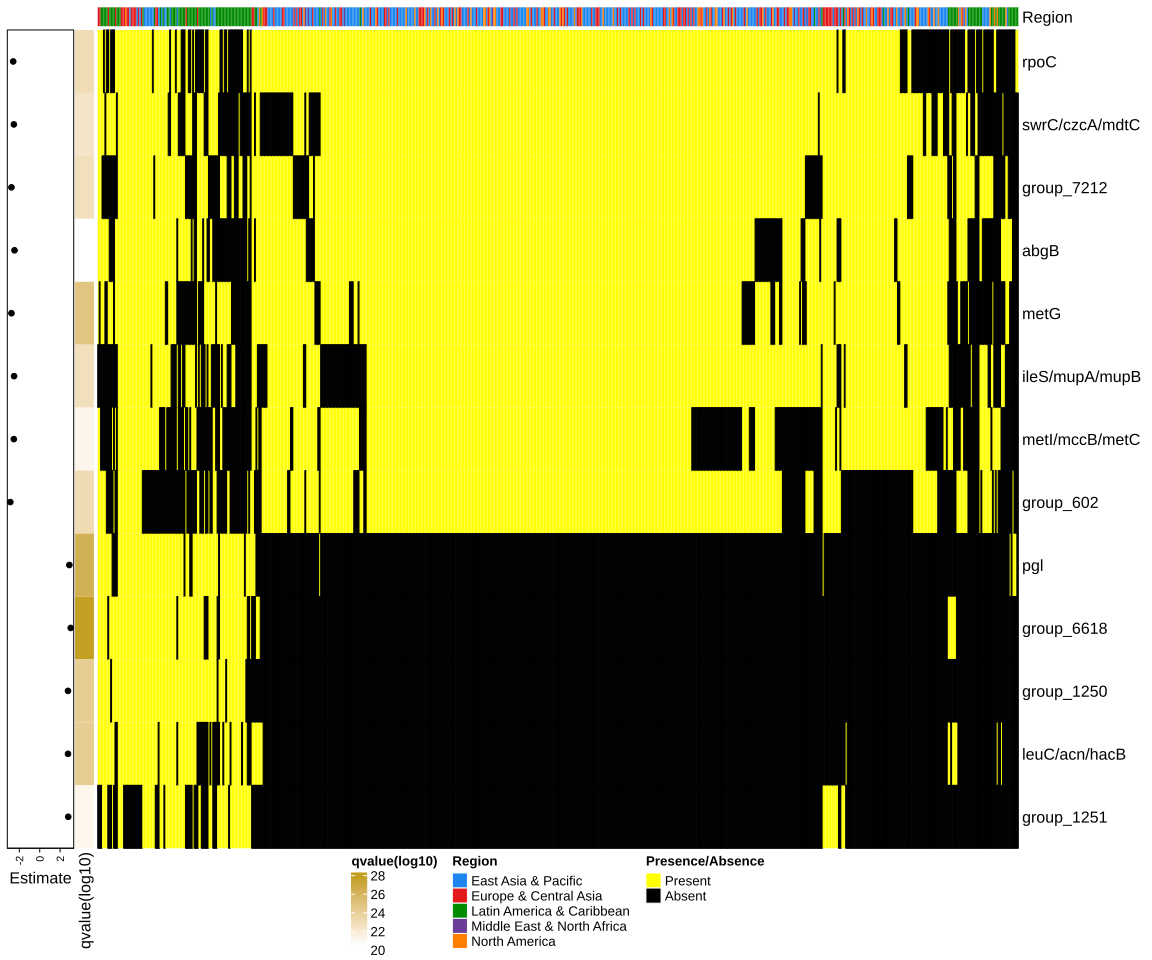

**Figure S10:** Heatmap showing the presence/absence pattern of differential present gene families across 640 MAGs annotated as *Aliscatomonas* sp900066535. Estimates and q-values were calculated by fitting a generalized linear model on each pangenome feature. P-values were corrected for multiple hypothesis testing using the FDR procedure. A positive estimate reflects whether the gene family was found to be more present in MAGs reconstructed from Honduras (top green bar annotation).

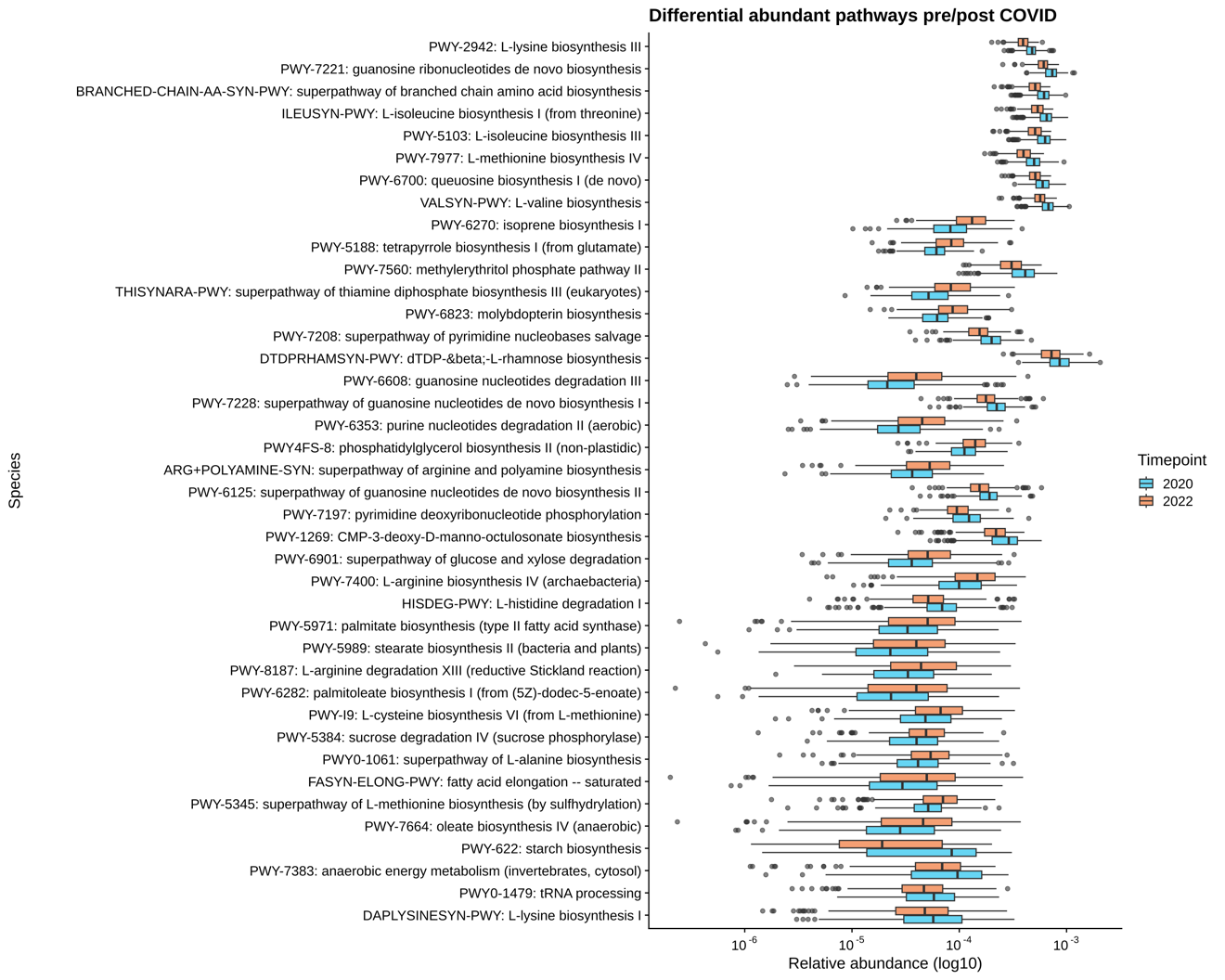

**Figure S11**: Top 40 differential abundant pathways identified in the 264 individuals positive for the presence of IgG only. Linear mixed-models as implemented by MaAsLin 2 identified a total of 248 MetaCyc pathways with relative abundance change in 2022 after Sars-CoV-2 infection.

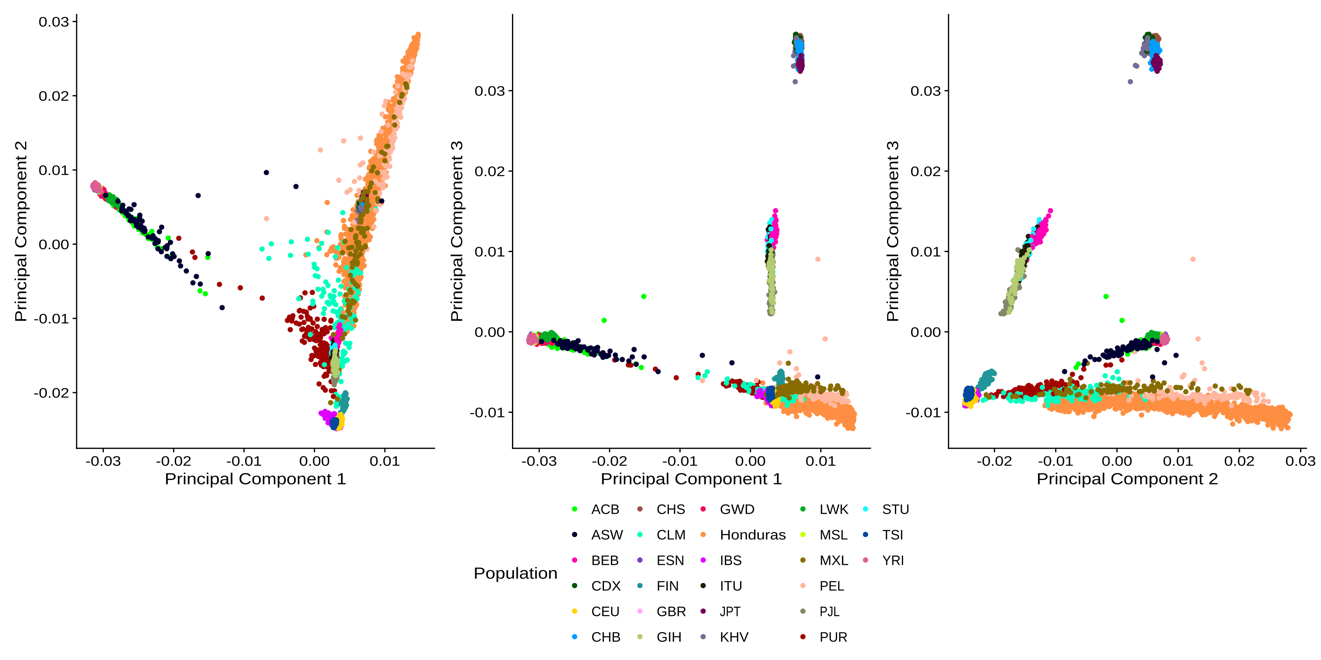

**Figure S12**: Principal Component Analysis of the 1,701 Honduras genomes showed alongside the 26 populations described in the 1000 Genomes project.

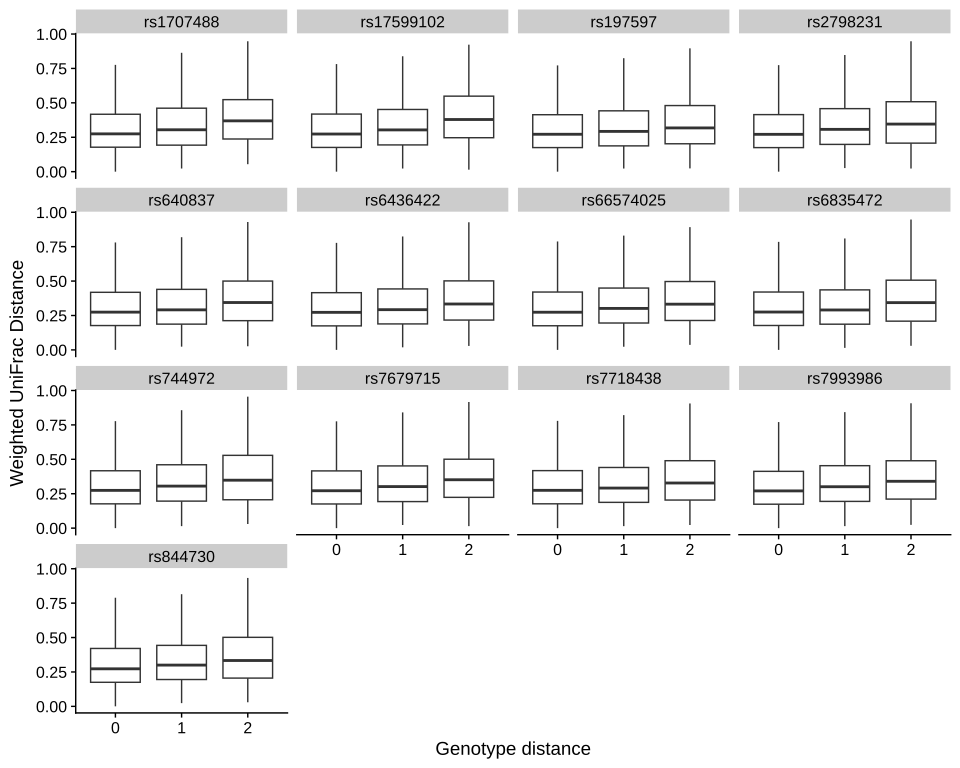

**Figure S13**: Distribution of microbiome distances calculated with the Weighted UniFrac distance are correlated with genetic distances calculated for the 13 SNVs identified as statistically significant by microbiomeGWAS. Genetic distances are calculated between pairs of alleles dosages.

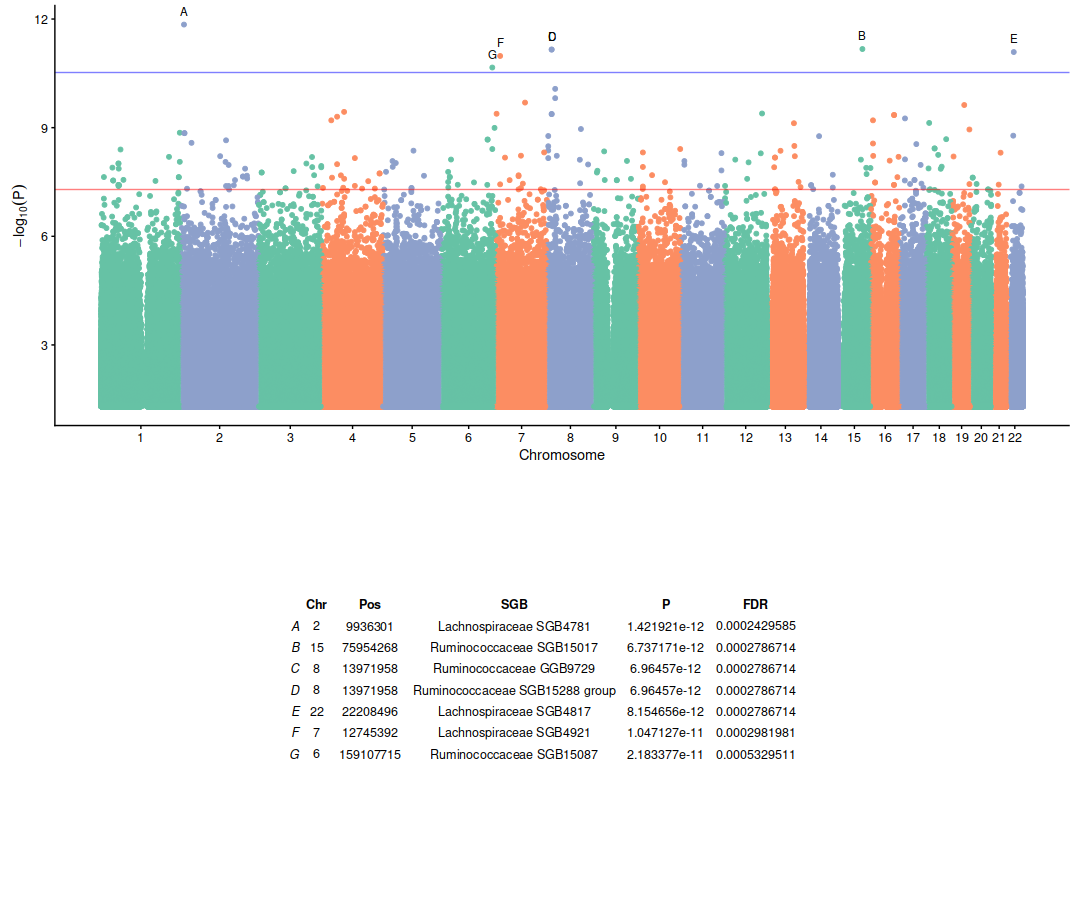

**Figure S14**: Manhattan plot of the eQTL analysis results. eQTLs showed are obtained by using species- and genus- level abundances as traits. Genome-wide significance (p = 5 × 10^-8^) and study-wide significance (p = 2.93 × 10^-11^) are reported as a red or blue lines. eQTL passing the study-wide significance threshold are annotated with a letter and their details are reported in the bottom table.

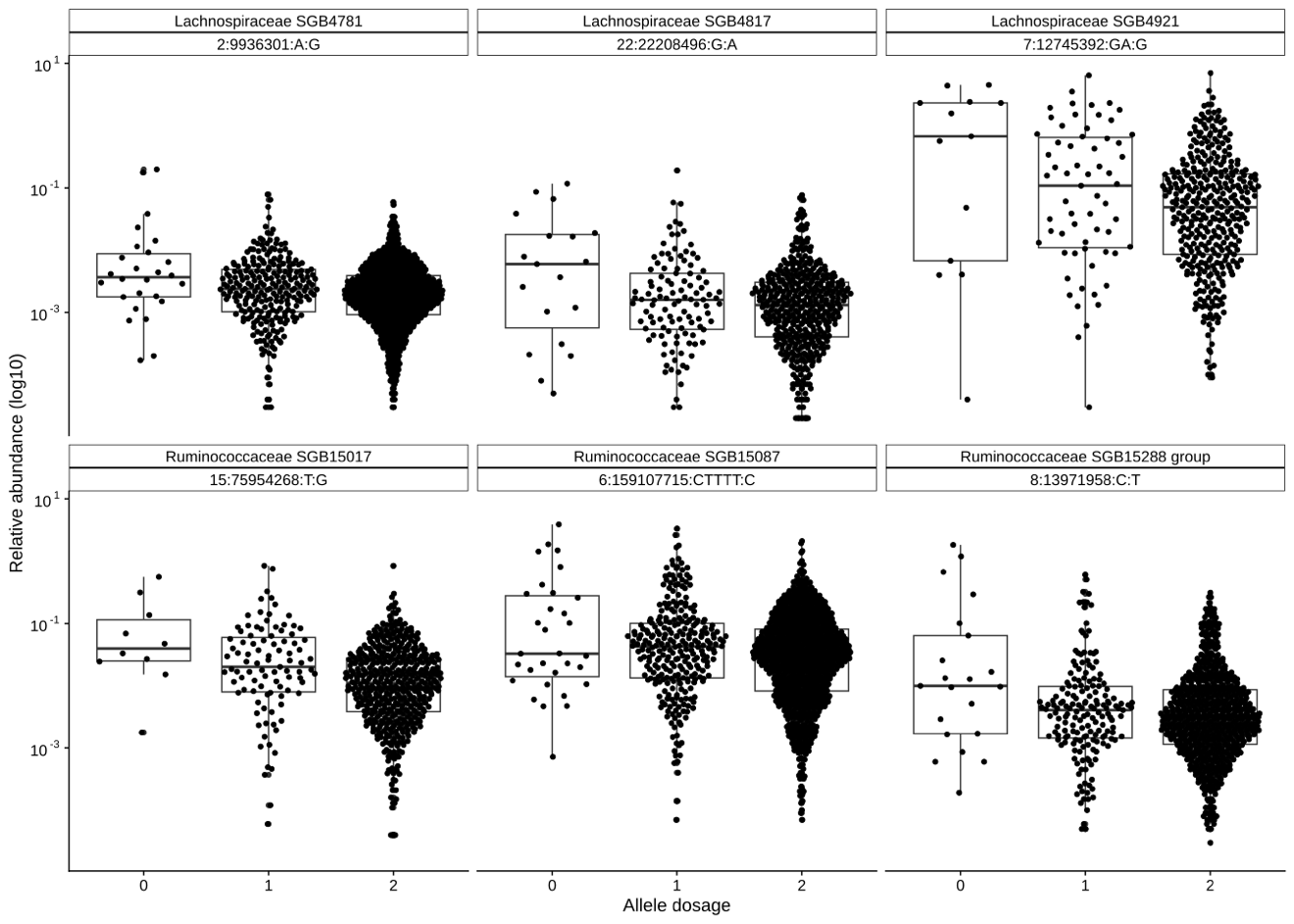

**Figure S15**: Distribution of relative abundances between different allele dosages for the 6 species-level eQTL identified significant at study-wide threshold.
